# An *Arabidopsis* long noncoding RNA modulates the transcriptome through interactions with a network of splicing factors

**DOI:** 10.1101/2020.01.03.894055

**Authors:** Richard Rigo, Jérémie Bazin, Natali Romero-Barrios, Michaël Moison, Leandro Lucero, Aurélie Christ, Moussa Benhamed, Thomas Blein, Stéphanie Huguet, Céline Charon, Martin Crespi, Federico Ariel

**Affiliations:** Institute of Plant Sciences Paris-Saclay (IPS2), CNRS, INRA, Universities Paris-Sud, Evry and Paris-Diderot, Sorbonne Paris-Cite, University of Paris-Saclay, Batiment 630, 91405 Orsay, France; Instituto de Agrobiotecnologia del Litoral, CONICET, FBCB, Universidad Nacional del Litoral, Colectora Ruta Nacional 168 km 0, 3000, Santa Fe, Argentina

**Keywords:** core splicing factors, flagellin, long noncoding RNA, PRP8a, SmD1b

## Abstract

Alternative splicing (AS) is a major source of transcriptome and proteome diversity in higher organisms. Long noncoding RNAs (lncRNAs) have emerged as regulators of AS through a range of molecular mechanisms. In *Arabidopsis thaliana*, the AS regulators NSRa and b, which affect auxin-driven lateral root formation, can interact with the *ALTERNATIVE SPLICING COMPETITOR (ASCO)* lncRNA. Here, we analyzed the effect of the knockdown and overexpression of *ASCO* at genome-wide level and found a high number of deregulated and differentially spliced genes, related to flagellin responses and biotic stress. In agreement, roots from *ASCO*-knocked down plants are more sensitive to flagellin. Surprisingly, only a minor subset of genes overlapped with the AS defects of the *nsra/b* double mutant. Using biotin-labelled oligonucleotides for RNA-mediated ribonucleoprotein purification, we found that *ASCO* binds to the highly conserved core spliceosome component PRP8a. *ASCO* deregulation impairs the recognition of specific flagellin-related transcripts by PRP8a and SmD1b, another spliceosome component, suggesting that *ASCO* function regulates AS through the interaction with multiple splicing factors. Hence, lncRNAs may interact in a dynamic network with many splicing factors to modulate transcriptome reprogramming in eukaryotes.

## Introduction

Alternative splicing (AS) of pre-mRNAs represents a major mechanism boosting eukaryotic transcriptome and proteome complexity (Chaudhary *et al*, 2019). In recent years, the advent of novel sequencing technologies allowed us to analyze entire genomes and complete pools of transcripts, leading to the identification of a wide variety of mRNA isoforms in higher organisms. More than 90% of intron-containing genes in humans and over 60% in plants are alternatively spliced (Pan *et al*, 2008; Wang *et al*, 2008; Marquez *et al*, 2012; Gerstein *et al*, 2014). The significant diversity in the number of transcripts compared to the number of genes suggests that a complex regulation occurs at transcriptional and post-transcriptional levels (Syed *et al*, 2012). Many mRNA isoforms derived from the same DNA locus are tissue-specific or are accumulated under particular conditions (Djebali *et al*, 2012). In humans, numerous studies suggest that the misregulation of RNA splicing is associated to several diseases (Boon *et al*, 2007; Tanackovic *et al*, 2011; Yoshida *et al*, 2011; Faial, 2015). In plants, AS plays an important role in the control of gene expression for an adequate response of plants to stress conditions (Palusa *et al*, 2007; Tanabe *et al*, 2007; Filichkin *et al*, 2010; Yan *et al*, 2012; Reddy *et al*, 2013; Ding *et al*, 2014; Zhan *et al*, 2015; Laloum *et al*, 2018; Jabre *et al*, 2019; Rigo *et al*, 2019). Alternative splicing modulates gene expression mainly by (i) increasing gene-coding capacity, thus proteome complexity, through the generation of a subset of mRNA isoforms derived from a single locus, and/or (ii) triggering mRNA degradation through the introduction of a premature termination codon in specific isoforms that would lead to nonsense mediated decay (NMD).

Besides the identification of an increasing number of AS events on mRNAs, next generation sequencing technologies led to the identification of thousands of RNAs with no or low coding potential (the so-called noncoding RNAs, ncRNAs), which are classified by their size and location respect to coding genes (Ariel *et al*, 2015). The long ncRNAs (lncRNAs, over 200 nt) act directly in a long form or may lead to the production of small ncRNAs (smRNAs) acting through base pairing recognition of their mRNA targets. There is growing evidence that large amounts of lncRNAs accumulate in particular developmental conditions or during diseases, suggesting that they participate in a wide range of biological processes. In recent years, several lncRNAs from higher organisms have been characterized as modulators of virtually every step of gene expression through interaction with proteins involved in chromatin remodeling, transcriptional control, co- and post-transcriptional regulation, miRNA processing, and protein stability during various developmental processes (Zhang *et al*, 2013; Ariel *et al*, 2015; Song *et al*, 2016). In particular, a growing number of lncRNAs have been linked to the modulation of AS in both plants and animals (Romero-Barrios *et al*, 2018). The main mechanisms involving lncRNAs in AS modulation have been classified as following: (i) lncRNAs interacting with splicing factors (Bardou *et al*, 2014; Barry *et al*, 2014; Ji *et al*, 2014; West *et al*, 2014; Kong *et al*, 2016); (ii) lncRNAs forming RNA-RNA duplexes with pre-mRNA molecules (Beltran *et al*, 2008; Villamizar *et al*, 2016) and (iii) lncRNAs affecting chromatin remodelling of alternatively spliced target genes (Gonzalez *et al*, 2015; Conn *et al*, 2017).

In *Arabidopsis thaliana,* the IncRNA *ASCO (ALTERNATIVE SPLICING COMPETITOR)* is recognized *in vivo* by the plant-specific NUCLEAR SPECKLE RNA-BINDING PROTEINS (NSRs), involved in splicing (Bardou *et al*, 2014). Notably, there is no evidence that ASCO undergoes splicing, although it is recognized by splicing factors. The analysis of a transcriptomic dataset of the *nsra/b* mutant compared to wild type (WT) plants, revealed an important number of intron retention events and differential 5’ start or 3’ end in a subset of genes, notably in response to auxin (Tran *et al*, 2016). Indeed, the *nsra/b* mutant exhibits diminished auxin sensitivity, e.g. lower lateral root (LR) number than WT plants in response to auxin treatment. This phenotype was related to the one observed for *ASCO* overexpressing lines. Interestingly, the splicing of a high number of auxin-related genes was perturbed in *nsra/b* mutants and several of them behaved accordingly in the *ASCO* overexpressing lines. The *ASCO*-NSR interaction was then proposed to regulate AS during auxin responses in roots (Bardou *et al*, 2014). More recently, an RNA-immunoprecipitation assay followed by RNA-seq (RIP-Seq) served to identify genome-wide RNAs bound *in vivo* by NSRa (Bazin *et al*, 2018). Long ncRNAs transpired to be privileged direct targets of NSRs in addition to specific NSR-dependent alternatively spliced mRNAs, suggesting that other lncRNAs than *ASCO* may interact with NSRs to modulate AS (Bazin *et al*, 2018).

In this work, we deeply characterize *ASCO* knocked-down plants and uncovered its general role in AS regulation, not only in response to auxin treatment. A transcriptomic analysis of *ASCO* knocked-down seedlings revealed a de-regulation of immune response genes and, accordingly, *ASCO* RNAi-silenced plants exhibited enhanced root growth sensitivity to flagellin 22 (flg22). The transcriptomic analysis of the *ASCO* over-expressing vs. *ASCO* knocked-down seedlings revealed distinct and overlapped effects on the whole of the mRNA population. Then, we assessed the genome-wide impact of *ASCO* function on AS and found many flg22-response regulatory genes showing differential AS in *ASCO*-deregulated lines. Surprisingly, the effect of *ASCO* knock-down on AS was clearly distinct from the defects of the *nsra/b* double mutant suggesting that *ASCO* impacts AS through a different interaction with the splicing machinery. Searching for *ASCO*-interacting proteins, we found SmD1b and PRP8a, two core components of the spliceosome that also recognize subsets of AS-regulated flg22-regulatory genes, also differentially spliced in *prp8a* (*prp8-7* allele; Sasaki *et al*, 2015) and *smd1b* mutants. Furthermore, *ASCO* overexpression competes for PRP8a and SmD1b binding to particular mRNA targets. Hence, lncRNAs may interact with key conserved components of the spliceosome to integrate a dynamic splicing network that modulates transcriptome diversity in eukaryotes.

## Results

### The *ASCO* lncRNA participates in lateral root formation

It was previously shown that *ASCO* (AT1G67105) overexpression results in a lower number of LRs in response to auxin treatment, a phenotype related to that of the *nsra/b* mutant, suggesting that increasing *ASCO* expression may lead to a titration of NSR activity in splicing (Bardou *et al*, 2014). To understand the role of *ASCO* in plant development, we generated independent RNAi lines to down regulate the levels of *ASCO* expression (RNAi*-ASCO1* and RNAi*-ASCO2*, Fig EV1A-B). Under control growth conditions, only RNAi-*ASCO1* plants exhibited a significantly longer primary root than the Col-0 wild type (WT) plants (Fig EV1C), whereas both independent lines showed an enhanced LR density in response to auxin treatment (Fig EV1D), the opposite phenotype to the one displayed by the *ASCO* overexpressing lines (Bardou *et al*, 2014). Furthermore, we transformed *A. thaliana* with a construct bearing 2631 bp of the *ASCO* promoter region controlling the expression of the fusion reporter genes *GFP*::*GUS* (*proASCO::GUS*). The *proASCO* construct includes the full intergenic region upstream of *ASCO* fusing the reporter gene to the first ATG in the *ASCO* locus at the position +15 from the transcription start site (Fig EV1E). In roots, *proASCO::GUS* was active very early in LR development, in pericycle cells undergoing the first division (Fig EV1F), whereas activity was then restricted to the vasculature adjacent to the LR primordium between stages II and VIII of LR development (according to Malamy and Benfey, 1997). Thus, *ASCO* expression pattern is in agreement with the LR-related phenotype of RNAi-*ASCO* plants.

### Deregulation of *ASCO* expression triggers a transcriptional response to biotic stress

In order to decipher the role of *ASCO* in the regulation of gene expression at a genome-wide level, we performed an RNA-seq of *A. thaliana* 14-day-old seedlings RNAi-*ASCO1* vs. WT Columbia (Col-0) accession in standard growth conditions. Overall, more genes were upregulated (321) than downregulated (178) in *ASCO* silenced plants (Fig 1A). Over 90% of deregulated transcripts correspond to protein-coding genes, according to Araport11 gene annotation (Fig 1B; Table EV1). To extend our understanding on the genome-wide role of *ASCO* in the control of gene expression, we searched the putative function of differentially expressed genes using Gene Ontology (GO). This analysis revealed a clear enrichment of deregulated genes involved in immune and defense response (FDR < 8e-4), as well as related pathways such as “response to chitin” and “glucosinolate metabolic pathways” (Fig 1C). Interestingly, related pathways were also partially observed in *nsra/b* mutants in response to auxin (Bazin *et al*, 2018). The upregulation of biotic stress-related genes was validated by RT-qPCR in both *RNAi-ASCO* lines compared to WT for a subset of 6 chosen transcription factors (TFs) which have been linked to the response to pathogens (Fig EV1G): *STZ* or *ZAT10* (AT1G27730) encodes for a Zn-finger TF involved in the response to oxidative stress (Munekage *et al*, 2015) and acts as a negative regulator of methyl jasmonate (MeJA) biosynthesis (Pauwels *et al*, 2008); MYB29 (AT5G07690) positively regulates the biosynthesis of aliphatic glucosinolate (AGSL), an essential defense secondary metabolite in *A. thaliana* (Li *et al*, 2013); WRKY33 (AT2G38470) controls the ABA biosynthetic pathway in response to the necrotrophic fungi *Botrytis cinerea* (Liu *et al*, 2015); ERF6 (AT4G17490) is a positive regulator of the MeJA and ethylene-mediated defense against *B. cinerea* (Moffat *et al*, 2012); ERF104 (AT5G61600) participates in the ethylene-dependent response to flg22 (Bethke *et al*, 2009); and ERF105 (AT5G51190) was shown to be strongly regulated in response to chitin (Libault *et al*, 2007) and to bind to the GCC-box pathogenesis-related promoter element (O’Malley *et al*, 2016). Remarkably, all these pathogen-related TFs are transcriptionally over accumulated in control conditions in the RNAi*-ASCO* plants (Fig EV1G), indicating that the deregulation of *ASCO* expression triggers molecular defense responses likely through the induction of pathogen-related TFs.

**Figure 1.**
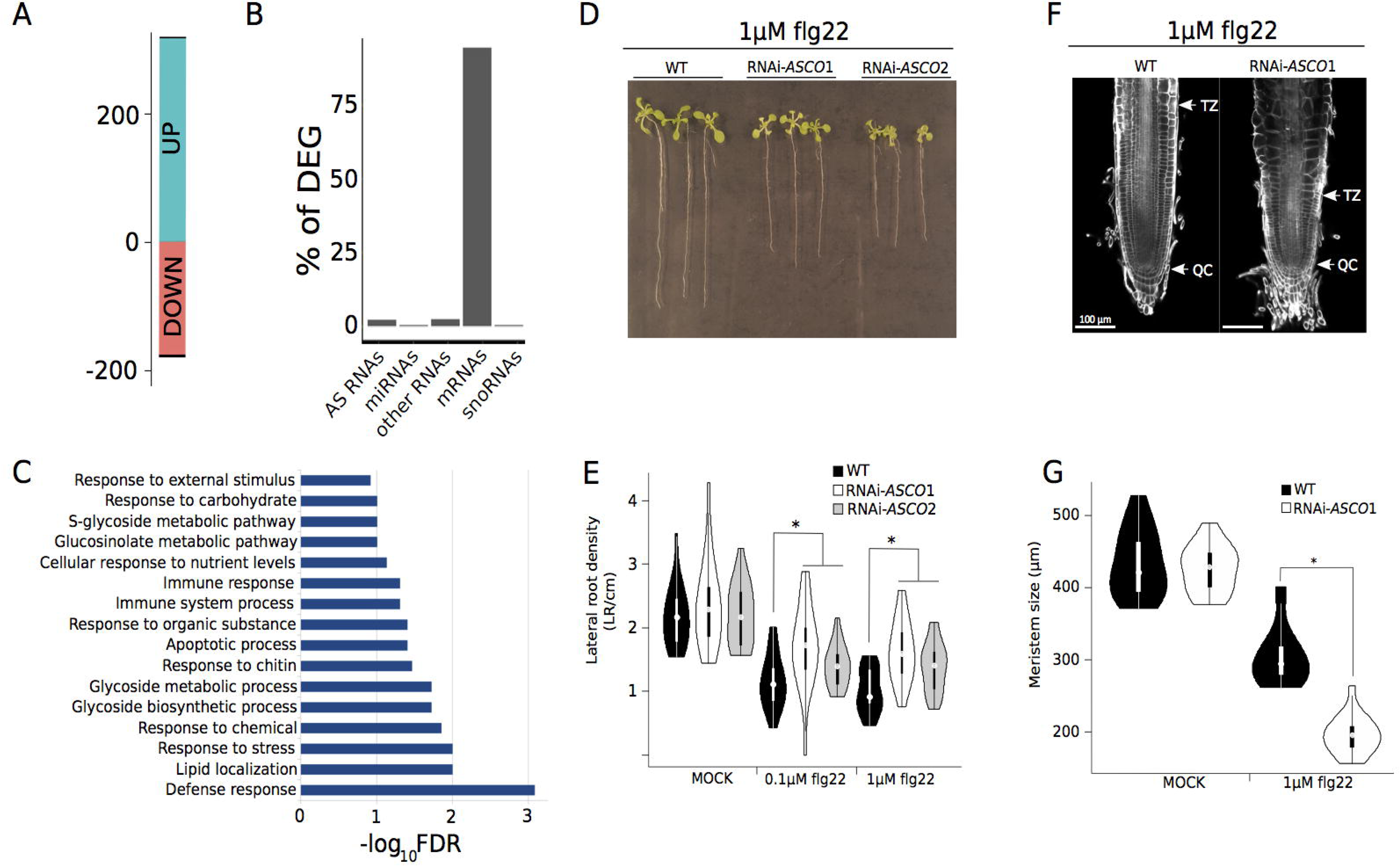
*ASCO* modulates steady-state levels of transcripts involved in plant immune responses affecting the sensitivity to flg22 peptide. **A** Number of differentially up- and down-regulated genes (DEG) in RNAi-*ASCO*1 seedlings as compared to wild type (WT) according to the RNA-seq approach (FDR < 0.01, |log2FC| >= 0.75). **B** Fraction of DEG found in each transcript class as defined in the Araport11 gene annotation. AS stands for antisense. **C** Enriched Gene Ontology (GO) of DEG in RNAi-*ASCO1* seedlings as compared to WT, x-axis represent the -log10 FDR for the enrichment of each GO category over genome frequency. **D** A representative picture of 14-day-old plants grown 9 days in liquid 1/2MS supplemented with 1 μΜ flg22. **E** Lateral root density of WT and two independent RNAi-*ASCO* lines 9 days after transfer in 1/2MS supplemented with 0.1 μΜ or 1 μΜ flg22. **F** A representative picture of root apical meristems after cell wall staining, in response to flg22. TZ: Transition Zone; QC: Quiescent Center. **G** Root apical meristem size of WT and RNAi-*ASCO1*. Error bars indicate the standard error. The asterisk (*) indicates a significant difference as determined by Mann Whitney’s U test p < 0.05, n = 18.

It is known that peptides corresponding to the most conserved domains of eubacterial flagellins (flg) act as potent elicitors in *A. thaliana*. Notably, flg22 causes callose deposition, induction of genes encoding for pathogenesis-related proteins and a strong inhibition of growth, including root development (Gomez-Gomez *et al*, 1999; Beck *et al*, 2014; Poncini *et al*, 2017). Thus, we first assessed the transcriptional behavior over time of *ASCO* in response to flg22. As shown in Fig EV1H, *ASCO* accumulation in roots was not significantly affected by flg22. Then, we characterized the physiological response of both *ASCO* RNAi-silenced lines to the exogenous treatment with flg22. Five-day-old plantlets were treated or not for 9 additional days with 0.1 or 1 μΜ flg22. Strikingly, the roots of RNAM*SC07* and *2* plants exhibited a normal development in control conditions (Fig EV3), whereas they were more sensitive to flg22 treatment, exhibiting a significantly shorter primary root (Figs 1D and 1F) but a minor reduction in the number of total LR, resulting in a higher density of LRs (Fig 1G). Cell wall staining and microscopic observation allowed us to quantify meristem size and determine that RNAi*-ASC0* plants show a reduction of the meristematic zone in response to flg22, e.g. shorter distance between the quiescent center and the beginning of the transition zone (Figs 1E, 1H and EV3B). This reduction in size is the result of a significant lower number of cells forming the root meristematic zone (Fig EV3C). Together with the physiological phenotype, we characterized the molecular response to this elicitor. A small subset of flg22-responsive genes was chosen (Asai *et al*, 2002; Zipfel *et al*, 2004; Boudsocq *et al*, 2010) to assess putative expression changes due to *ASC0* knock-down. In mock conditions, RNAi-*ASC0* lines exhibited an increased expression for certain flg22-responsive genes tested (Fig EV3D). Interestingly, this subset of genes suffered an overall lower induction by 3h flg22 in RNAi-*ASC0* plants (Fig EV3E), in agreement with the previously observed altered sensitivity to flg22 of RNAi-*ASC0* roots.

To further demonstrate the link between *ASC0* and the response to flg22, we searched for additional independent Arabidopsis lines exhibiting a deregulation in *ASC0* accumulation. We characterized two insertional mutants located at the 5’ region (*asco*-1) and the 3’ region of the locus (*asco*-2; Fig EV4A). The first line, *asco-1,* resulted to be an over-expressor of a truncated *ASC0* version (lacking a minor portion of the 5’ region) whereas the *asco-2* T-DNA line shows minor changes in *ÆSC0* expression (Fig EV4B). Interestingly, the nearly 50-fold over-accumulation of *ASC0* RNA in *asco-1* plants results in a slight reduction of LR density in response to flg22, opposite to RNAi lines (Fig EV4E). Accordingly, when we assessed two independent *35S:ASC0* over-expressing lines, reaching an over-accumulation of 1000- to 2500-fold RNA levels, plants exhibit a longer main root and a lower density of LRs in response to fg22 (Fig. 4AB). Therefore, *ÆSC0* participates in the regulation of biotic-stress related genes, shaping root architecture in response to flg22.

### *ASCO* modulates the alternative splicing but not the accumulation of a subset of pathogen-related mRNAs

Considering that overexpression of *ASC0* affected the AS of NSR mRNA targets (Bardou *et al*, 2014), we searched for mis-spliced genes potentially explaining the global physiological impact of *ASCO* deregulation. To this end, we used two complementary approaches to detect both differential AS based on annotated isoforms (Reference Transcript Dataset for *A. thaliana*, AtRTD2; Zhang *et al*, 2017), and differential RNA processing using RNAprof (Tran *et al*, 2016). Based on RNAprof, a total of 303 differential RNA processing events in 281 distinct genes were identified comparing RNAM*SCO* with WT plants in control growth conditions (Table EV2), whereas the SUPPA method (Trincado *et al*, 2018) identified 205 genes with evidence of differential AS in the AtRTD2 database. Comparison of the two analyses with differentially expressed genes (DEG) in RNAi-*ASCO* lines revealed that most differentially alternatively spliced (DAS) genes are not differentially accumulated (Fig 2A). In addition, our analyses showed the complementarity between the two approaches since only 24 common DAS genes were identified by both methods. Classification of the location and the relative isoform accumulation (up or down) of these events revealed that the majority of them were located in introns and had higher read coverage in RNAi*-ASCO* plants, suggesting that *ASCO* inhibited proper intron splicing on these genes (Fig 2B). Nevertheless, we also identified differential events located within 5’UTR, CDS or 3’UTR suggesting that other RNA processing events, in addition to intron retention, are affected by *ASCO* expression levels. Analysis of differential AS events with SUPPA revealed 317 significant DAS events (|dPSI| > 0.1, pval < 0.01) on 205 unique genes from the AtRTD2 transcript annotation database (Table EV3). Similarly to the analysis with RNAprof, most of these events corresponded to retained introns (62%) but we also identified a significant number of alternative 3’ splice site and alternative 5’ splice site selection modulated in *ÆSCO* knock-down lines (Fig 2C). To determine the most significant impact of the AS events, we sought to identify isoform switching events (where major changes in a specific isoform leads to a significant change on protein composition) using the IsoformSwitchAnalyzeR package (Table EV4; Vitting-Seerup and Sandelin, 2017). Strikingly, isoforms switching events were detected for 52 genes, out of which 12 and 34 were common cases detected by RNAprof and AtRTD2-SUPPA, respectively (Fig 2D). *In silico* analysis of switching isoforms protein sequences identified that the AS events may lead to (i) a change of ORF length; (ii) gain or loss of conserved PFAM protein domain and signal peptides; (iii) or a change of the coding potential and the sensitivity to NMD (Fig 2E). Since AS can often trigger NMD, an important mechanism of plant gene expression regulation (Kalyna *et al*, 2012), we compared DAS genes to those transcripts over-accumulated in the double mutant of the NMD factor homologs *UP FRAMESHIFT1* (*UPF1*) and *UPF3*, *upf1-upf3* (Drechsel *et al*, 2013). As shown in Fig 2F, 66 and 29 genes were reported by RNAprof and SUPPA, respectively. Hence, the majority of AS events controlled by *ASCO* seem to be independent of the UPF1-UPF3 mediated RNA quality control machinery, at least in the conditions previously assessed.

**Figure 2.**
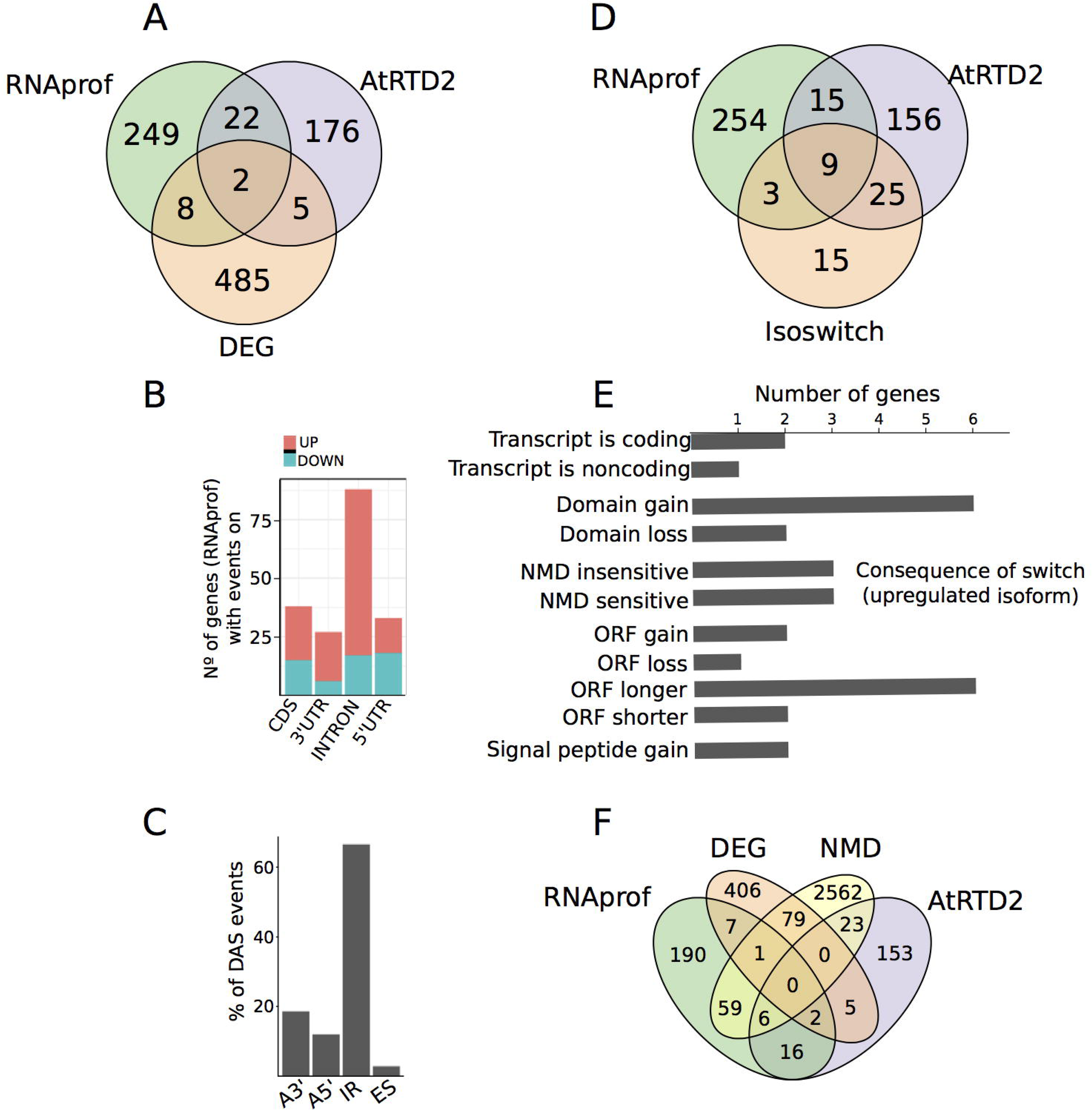
*ASCO* modulates alternative splicing. **A** Comparison of differentially processed transcripts (RNAprof) and differential AS genes (AtRTD2-SUPPA) with Differentially Expressed Genes (DEG). **B** Number of genes containing at least one differential RNA processing event (as defined by RNAprof p.adj < 0.001) in CDS, introns, 5’UTR and 3’UTR between RNAi-*ASCO1* and WT. Up and Down fractions correspond to increase or decrease, respectively, of RNA-seq coverage in RNAi-*ASCO1* for each specified gene feature. **C** Proportion of DAS events identified by AtRTD2-SUPPA in RNAi-*ASCO1* compared to WT; Alternative 3’ site (A3’), Alternative 5’ site (A5’), Intron Retention (IR), Exon skipping (ES). **D** Comparison of differentially processed transcripts (RNAprof) and differentially AS genes (AtRTD2-SUPPA) with genes showing significant isoform switch events (Isoswitch). **E** Summary of the predicted consequence of the isoform switch events as shown by the feature acquired by the upregulated isoform. ncRNA stands for noncoding RNA, and NMD stands for nonsense mediated decay. **F** Comparison of differentially processed transcripts (RNAprof), differentially AS genes (AtRTD2-SUPPA) and differentially expressed genes (DEG) with genes significantly upregulated in the *upf1upf3* mutant, indicating genes potentially regulated by NMD.

Furthermore, we performed an RNA-seq of 14-day-old seedlings *35S:ASCO1* vs. Col-0 WT plants in standard growth conditions. Interestingly, there is a minimal overlap between DEG and DAS genes in WT vs. RNAi-*ASCO1* and *35S:ASCO1*. Strikingly, the up- and down-deregulation of *ASCO* resulted in alternative subsets of DAS genes, including only 120 common DAS between RNAi-*ASCO1* and *35S:ASCO1*, compared to 227 and 137 excluding events, respectively (Fig. 3C). Further comparison of DEG fold change revealed a global correlation of gene expression changes in 35S:*ASCO1* and RNAi-*ASCO1* as compared to WT but also that specific gene subset specifically responded to down or up-regulation of *ÆSCO* (Fig 3D).

**Figure 3.**
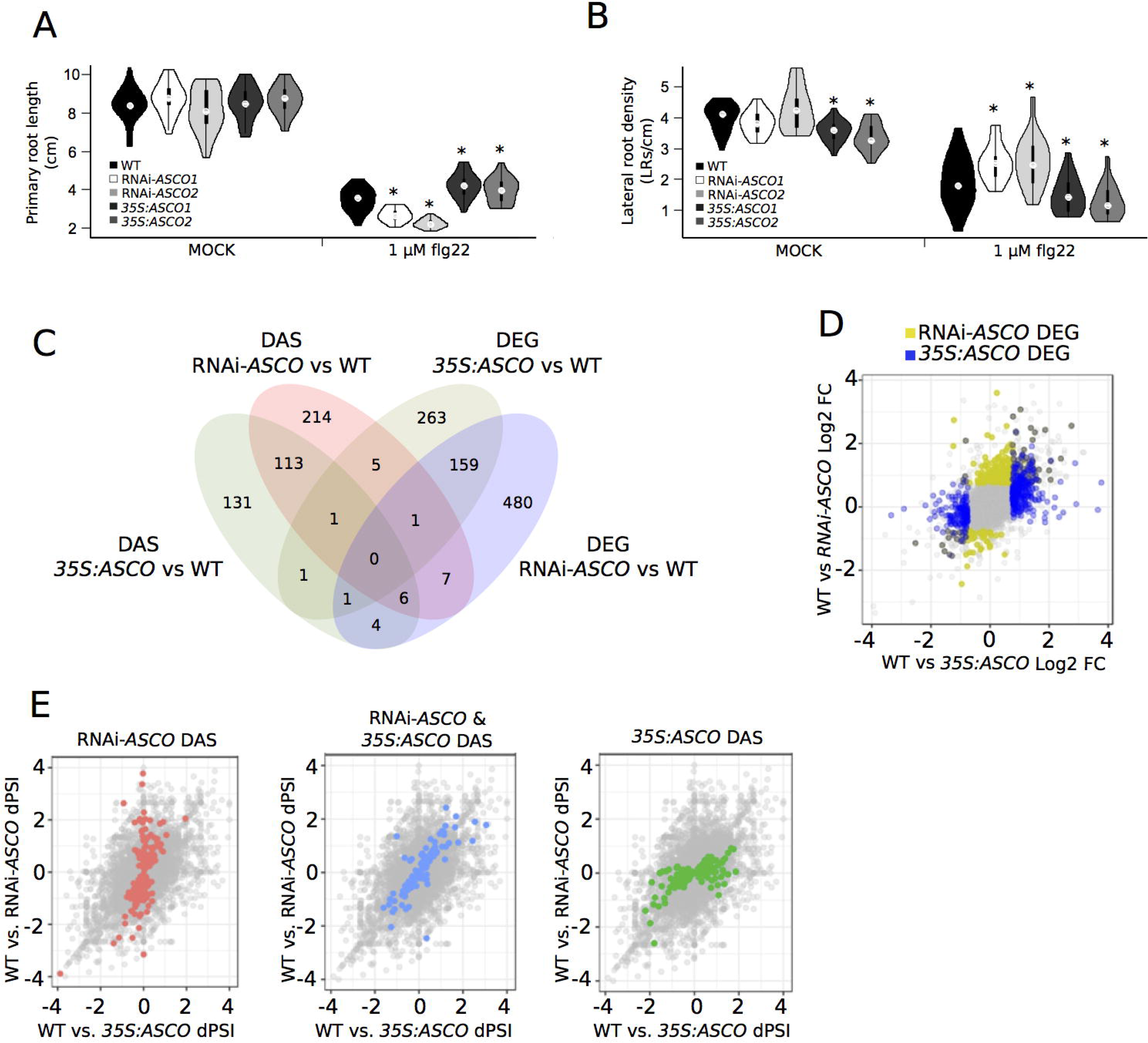
RNAi:ASCO and *35S:ASCO* lines exhibit altered sensitivity to flg22 peptide and share some DEG and DAS targets. **A, B** Primary root length (**A**) and lateral root density (**B**) of WT, two RNAi-*ASCO* and two *35S:ASCO* independent lines 9 days after transfer in 1/2MS supplemented with 1 μΜ flg22. Error bars indicate the standard error. The asterisk (*) indicates a significant difference as determined by Mann Whitney’s U test < 0.05, n = 24. **C** Overlap between differentially expressed (DEG) and spliced (DAS) genes in RNAi-*ASCO* and *35S:ASCO* as compared to WT. **D** Scatter plot showing the respective gene expression fold change in RNAi-*ASCO* and *35S:ASCO* lines as compared to WT. Genes showing significant changes in RNAi-*ASCO* and *35S:ASCO* are highlighted as yellow and blue dots, respectively. **E** Scatter plot showing the respective Percent Spliced In difference (dPSI) in RNAi-*ASC*O and *35S:ASCO* lines as compared to WT. Genes showing significant changes in RNAi-*ASCO*, *35S:ASCO* or in both lines are highlighted as red, green or blue dot, respectively.

Similarly we compared of dPSI (difference of Percent Spliced In), which represent the change of each AS event. The analysis revealed that the group of 120 common AS events are positively correlated between the two lines as compared to wild type (Fig 3E). In addition, this also revealed that dPSI of AS events significantly regulated in response to either overexpression or silencing of *ASCO,* respectively, were not correlated between the two lines (Fig 3E). Overall, the effect of *ASCO* silencing was more extensive on DAS, compared to its overexpression.

In order to better understand the impact of *ASCO* deregulation on the plant response to flg22, we focused on the transcriptional behavior of specific genes. Strikingly, several pathogen-related genes appeared differentially spliced in RNAi*-ASCO1* line although they were not affected in their global expression levels (Fig 2A). These AS events include two members from the NB-LRR disease resistance genes, *RPP4* (Mohr *et al*, 2010), as well as the splicing regulatory serine-rich protein coding gene *SR34* (AT1G02840), needed for accurate response to pathogens (Xu *et al*, 2011). The splicing of the *SR34* own pre-mRNA is auto-regulated and depends on the activity of immune response factors (Zhang *et al*, 2014). Other relevant AS targets are *SNC4* (AT1G66980) which encodes a receptor-like kinase that participates in the activation of the defense response and its AS is impaired in defense-related mutants, affecting the response to pathogens (Zhang *et al*, 2014); *SEN1* (AT4G35770), a senescence marker gene primarily regulated by salicylic acid (SA)- and MeJA-dependent signaling pathways (Schenk *et al*, 2005) and *NUDIX HYDROLASE7* (*NUDT7*; AT4G12720) which regulates defense and cell death against biotrophic pathogens (Straus *et al*, 2010). Another interesting target is *EPITHIOSPECIFIER PROTEIN* (*ESP*; AT1G54040) a gene involved in the plant defense to insects and also differentially spliced in response to MeJA (Kissen *et al*, 2012) although this gene was differentially expressed in RNAi-*ASCO* plants. We also included in the following analysis a NAD(P)-binding Rossmann-fold protein family Gene *(NRG)* AT2G29290, exhibiting drastically altered AS upon *ASCO* knock-down. Specific DAS events first identified *in silico* (Fig 4AD, EV5 and EV6) were validated by RT-PCR and polyacrylamide gel electrophoresis by calculating the ratio between alternatively spliced and fully spliced isoforms (Isoform ratio, Figs 4BCEF, and EV6). Some events were also validated by quantitative RT-qPCR where each differential event was normalized with respect to an internal gene probe (called INPUT) which corresponds to a common exon. This allowed to calculate the splicing index (defined in the method section,Figs EV7). Altogether, our results indicate that the knock-down of *ASCO* expression affects the AS of a subset of genes whose isoforms distribution may modulate the pathogen-related transcriptome and affect the response to flg22.

**Figure 4.**
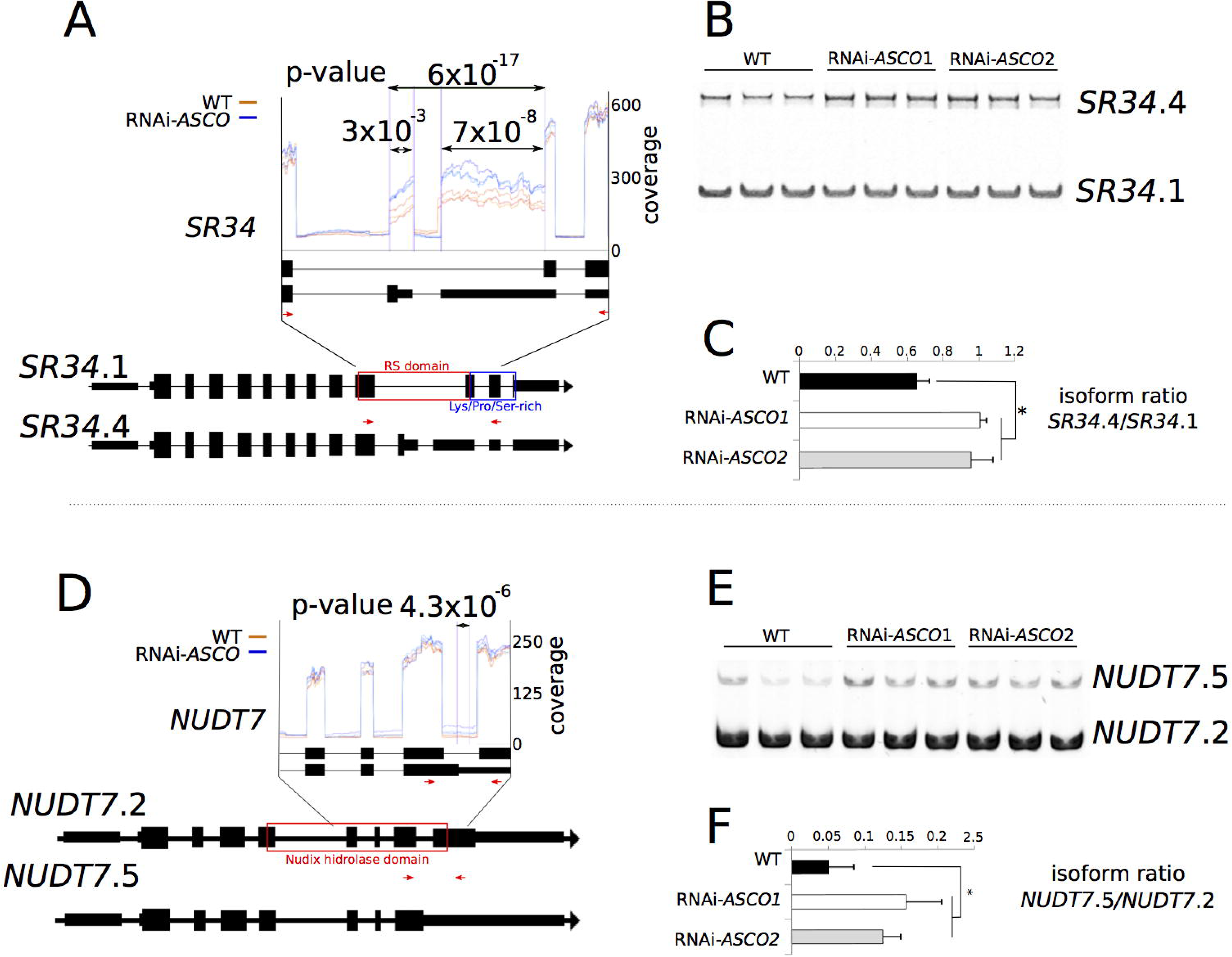
Validation of AS events in RNAi-*ASCO* lines. **A, D** Differential RNA processing events of *SR34* (AT1G02840) and *NUDT7* (AT4G12720) transcripts detected by RNAprof from the comparison of RNA-seq libraries of 14-day-old WT (red) and RNAi-*ASCO1* (blue) plants. Three biological replicates were used. Vertical purple lines and p-values indicate significant differential processing events. Structure of *SR34* (**A**) and *NUDT7* (**D**) RNA isoforms. Large black boxes indicate exons, narrow black boxes indicate UTRs, black lines indicate introns. Colored boxes indicate protein domains affected by an AS event. Red arrows indicate probes used for gel electrophoresis. **B, E** Analyses of RT-PCR products of *SR34* (**B**) and *NUDT7* (**E**) transcripts on 8% acrylamide gel. **C, F** Quantification of the ratio of *SR34* (**C**) and *NUDT7* (**D**) isoforms detected in the gel in (**B**) and (**E**) respectively. RNAs were extracted from WT and RNAi-*ASCO1* 14-day-old plants. The asterisk (*) indicates a significant difference as determined by Student’s T test < 0.05, n = 3.

### *ASCO* interacts with the spliceosome core components PRP8a and SmD1b

The fact that *ASCO* interacts with NSRs strongly suggested that *ASCO* would affect a large subset of NSR-targeted AS events. Surprisingly, DAS genes in the RNAi*-ASCO* and *35S:ASCO* plants only partially coincide with those in *nsra/b* double mutants (in response to auxin or not). In all, out of the 589 DAS events identified in *nsra/b* compared to WT, only 140 (32%) are common with RNAi*-ASCO1* and 109 (33%) with *35S:ASCO*, what represents 24% and 19% of all DAS events in the *nsra/b* mutant, respectively (Fig EV8A). Furthermore, *nsra/b* plants do not respond to flg22 in the same way as RNAi-*ASCO* plants (Fig EV8C and D), indicating that *ASCO* further modulates AS in an NSR-independent manner by a yet uncovered mechanism, notably affecting the response to biotic stress. Therefore, in order to decipher the AS-related complexes implicating *ASCO*, we performed an antisense oligonucleotides based pull-down method, related to the Chromatin Isolation by RNA Purification (ChIRP; Chu *et al*, 2011; Ariel *et al*, 2014) using nuclear extracts to purify *ASCO*-containing RNPs. Eighteen biotinylated probes matching *ASCO* were used in independent sets called EVEN and ODD, respectively (Table EV5). *ASCO*-ChiRP was performed in 4 biological replicates for each set of probes: ODD, EVEN, and an additional set matching the *LacZ* RNA, used as a negative control. *ASCO* enrichment was corroborated by ChiRP followed by RNA purification and RT-qPCR (Fig 5A). We then performed a mass spectrometry on the proteins from the purified *ASCO*-containing RNP to identify potential *ASCO* protein partners. Strikingly, among the RNA-related proteins identified in the EVEN and ODD samples, but not in the LacZ, we found the pre-mRNA-processing-splicing factor 8A, PRP8a. PRP8 is a core component of the spliceosome highly conserved in higher organisms and null mutants are generally embryo lethal (Grainger and Beggs, 2005). In *Arabidopsis*, a PRP8a leaky mutation was found to also affect the AS of the *COOLAIR* lncRNA (Marquardt *et al*, 2014) and results in a high number of intron retention events (Sasaki et al., 2015). Therefore, we developed specific antibodies against PRP8a and we tested them in immunolocalization experiments that revealed a nuclear localization pattern (Fig EV9) similar to what was previously observed in *Drosophila* (Claudius *et al*, 2014). In order to *in vivo* validate the interaction with *ASCO*, we performed an RNA-immunoprecipitation assay followed by qPCR (RIP-qPCR) from nuclear extracts, showing that PRP8a can recognize the spliceosomal *U5* RNA (Koncz *et al*, 2012) taken as a positive control, as well as the *ASCO* lncRNA (Fig 5B). The efficiency of the PRP8a immunoprecipitation (IP) was assessed by western blot comparing nuclei input samples, against the unbound fraction after IP, as well as the anti-PRP8a IP and the anti-IgG IP (Fig 5C). We then assessed the binding of PRP8a to the pathogen-related mRNAs differentially spliced in the RNAi*-ASCO* lines. PRP8a was indeed able to interact with 4 of these *ASCO*-related DAS genes. Furthermore, their binding was impaired upon the overexpression of *ASCO* (Fig 5D), hinting an *ASCO*-mediated competition of these mRNAs inside the PRP8a-containing spliceosome complex. Interestingly, *ASCO* is over-accumulated in the *prp8a* mutant allele (Sasaki *et al*, 2015 hereafter referred to as *prp8a*) background (Fig 6A), as it occurs in the *nsra/b* mutant plants (Bardou *et al*, 2014). Remarkably, similar AS defects were shown between the *prp8a* mutant and RNAi*-ASCO* lines for pathogen-related genes (Figs 6BC and EV10), indicating that the flg22 differential phenotype of RNAi*-ASCO* plants may be related to the interaction with the spliceosome components. Recently, we identified another core component of the spliceosome, SmD1b, linked to both AS and the recognition of aberrant ncRNAs to trigger gene silencing (Elvira-Matelot *et al*, 2016). Interestingly, *ASCO* expression levels are also increased in the *smd1b* mutant (Fig 6D), exhibiting the same transcript accumulation as in *prp8a* and *nsra/b* mutants. Hence, we wondered whether this other core component of the spliceosome also interacts with *ASCO* lncRNA. Using *pUBI*:*SmD1b-GFP* plants (*smd1b* background, Elvirat-Matelot., 2016), we performed a RIP assay and found that SmD1b also recognizes *ASCO in vivo* (Fig 6E) as well as the *U6* RNA taken as a positive control. Furthermore, SmD1b recognizes the four pathogen-related transcripts assessed (Fig 6F) although only two out of three pathogen-related transcripts were DAS in *smd1b* mutants: *ESP*, *SR34*, but not *NUDT7* (Fig 6GH, and EV10). *SNC4* total transcript levels were dramatically reduced in the *smd1b* mutant, hindering the analysis of relative isoforms accumulation (Fig EV10J). Altogether, our results indicate that *ASCO*, an apparently intron-less lncRNA, interacts with PRP8a and SmD1b, two core components of the spliceosome, contributing to determine the dynamic ratio between hundreds of alternatively spliced mRNAs, notably pathogen-related genes. Hence, lncRNA are able to interact with multiple core components of the splicing machinery to modulate the splicing patterns of particular subsets of mRNAs.

**Figure 5.**
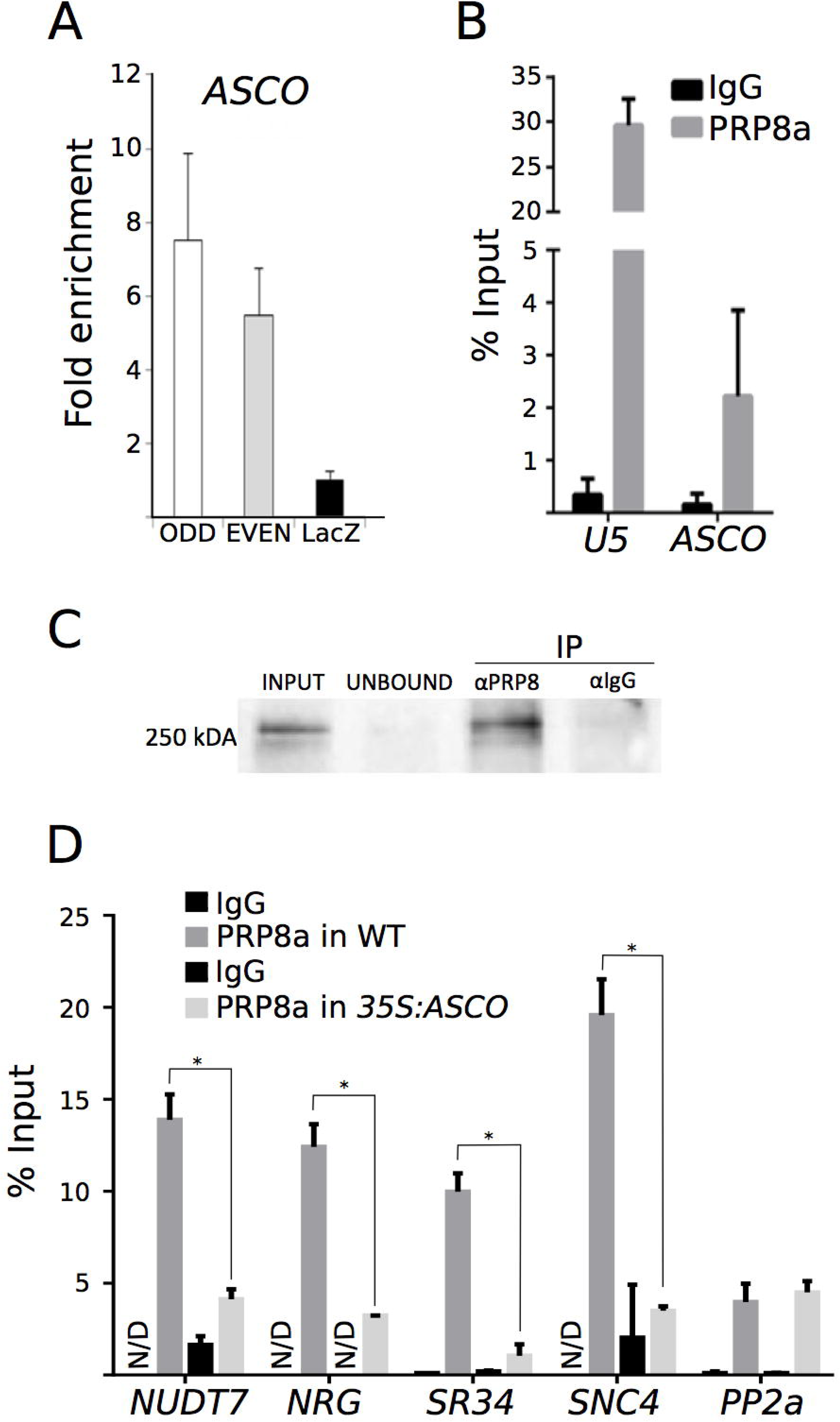
*ASCO* modulates the activity of the spliceosome component PRP8a. **A** Analysis of *ASCO* enrichment by ChIRP using two sets of independent biotinylated probes ODD and EVEN compared to negative control with probes designed against the *LacZ* RNA. The fold enrichment was calculated between ODD or EVEN samples against LacZ. These samples were used for protein precipitation and Mass Spectrometry analyses (ChIRP-MS), from which PRP8a was identified as a potential *ASCO* partner. **B** Validation of PRP8a-*ASCO* interaction by PRP8a-RIP. *U5* RNA was used as a positive control. C Immunoblot analysis was performed on input, unbound and IP fraction (anti-GFP and anti-IgG negative control) **D** PRP8a recognition of a subset of DAS RNAs is impaired in the *35S:ASCO* plants. In **B** and **D**, the results are expressed as a percentage of the Input for PRP8a RIP followed by RT-qPCR, and IgG RIP was used as a negative control. Error bars represent the standard error of 3 biological replicates. N/D stands for non-detectable.

**Figure 6.**
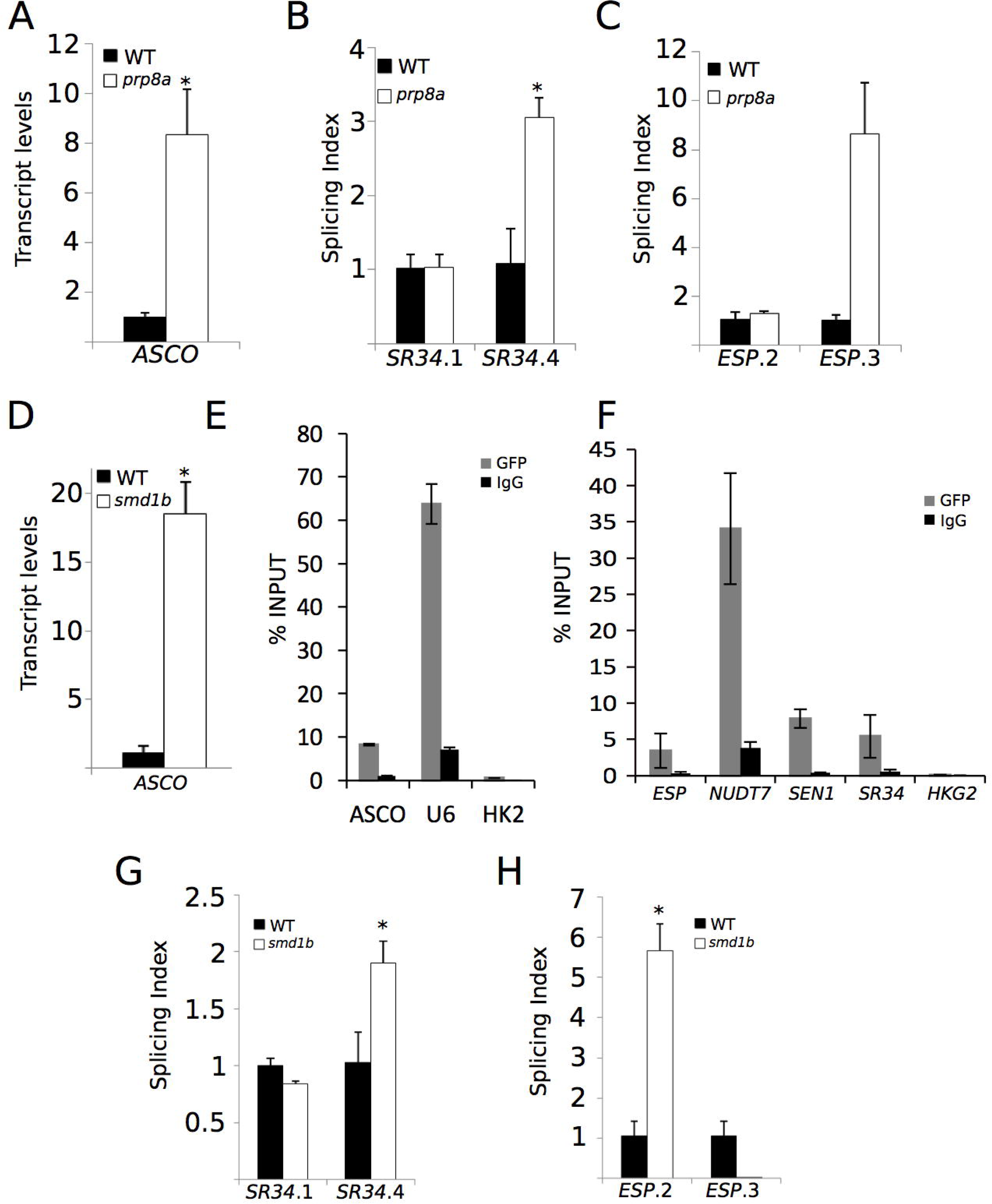
PRP8a and SmD1b regulate AS of *ASCO* mRNA targets. **A** *ASCO* transcript levels in WT and *prp8a* mutants. **B, C** *prp8a* leaky mutant displays similar AS events as observed in RNAi-*ASCO*. Quantification of *SR34* (**B**) and *ESP* (**C**) isoforms by RT-qPCR. **D** *ASCO* transcript levels in *smd1b* mutant. In **A** and **D** RNAs were extracted from WT, *prp8a* and *smd1b* 14-day-old plants. **E** SmD1b can bind *ASCO in vivo*. *U6* RNA was used as a positive control and a housekeeping gene (*HKG2*, AT4G26410) RNA as a negative control. **F** SmD1b recognizes *in vivo* the RNAs of 4 genes regulated by *ASCO*. In **E** and **F** the results were expressed as % INPUT in SmD1b-RIP and IgG-RIP used as a negative control. **G, H** *smd1b* mutant displays similar AS events as observed in RNAi-*ASCO*. Quantification of *SR34* (**G**) and *ESP* (**H**) isoforms by RT-qPCR. The asterisk (*) indicates a significant difference as determined by Student’s T test < 0.05, n = 3.

## Discussion

### Long noncoding RNAs modulate splicing regulatory networks

We show here that reducing *ASCO* expression has a major effect on AS at genome-wide level in plants. In animals, different splicing factors can recognize lncRNAs *in vivo* (Romero-Barrios *et al*, 2018), e.g. Y-BOX BINDING PROTEIN 1 (YBX1; Suresh *et al*, 2018), POLY(RC) BINDING PROTEINS 1 and 2 (PCBP1/2; van Dijk *et al*, 2015), FOX proteins (Yin *et al*, 2012) and the serine-rich splicing factors, such as SRSF6 (Kong *et al*, 2016), among others. The lncRNA *GOMAFU* is recognized through a tandem array of UACUAAC motifs by the splicing factor SF1, which participates in the early stages of spliceosome assembly (Tsuiji *et al*, 2011). Furthermore, *GOMAFU* was found to directly interact with the splicing factors QUAKING homolog QKI and SRSF1 (Barry *et al*, 2013). In adult mice, *GOMAFU* is expressed in a specific group of neurons and has been implicated in retinal cell development (Rapicavoli and Blackshaw, 2009; Rapicavoli *et al*, 2010), brain development (Mercer *et al*, 2010) and postmitotic neuronal function (Sone *et al*, 2007; Mercer *et al*, 2008). *GOMAFU*’s downregulation leads to aberrant AS patterns of typically schizophrenia-associated genes (Barry *et al*, 2013). Other lncRNAs recognized by splicing factors are *NUCLEAR PARASPECKLE ASSEMBLY TRANSCRIPT 1* (*NEAT1*) and *NEAT2* (also known as *METASTASIS ASSOCIATED LUNG ADENOCARCINOMA TRANSCRIPT 1; MALAT1)* (Hutchinson *et al*, 2007). RNA FISH analyses revealed an intimate association of *NEAT1* and *MALAT1* with the SC35 splicing factor containing nuclear speckles in both human and mouse cells, suggesting their participation in pre-mRNA splicing. Indeed, the *ASCO* lncRNA also interacts with NSRs and SmD1b both localized in nuclear speckles, whereas we show here that PRP8a seems to have nuclear localization in Arabidopsis. It was shown that *NEAT1* localizes to the speckles periphery, whereas *MALAT1* is part of the polyadenylated component of nuclear speckles (Hutchinson *et al*, 2007). *MALAT1* acts as an oncogene transcript and its aberrant expression is involved in the development and progression of many types of cancers (Wang *et al*, 2015; Li *et al*, 2017; Malakar *et al*, 2017). *MALAT1* can promote metastasis by interacting with the proline- and glutamine-rich splicing factor SFPQ, blocking its tumor suppression activity (Ji *et al*, 2014). In plants, little is known about the interaction between splicing factors and lncRNAs (Ariel *et al*, 2015; Romero-Barrios *et al*, 2018). NSRs are a family of RNA binding proteins that act as regulators of AS and auxin-regulated developmental processes such as lateral root formation in *A. thaliana*. These proteins were first shown to interact with some of their alternatively spliced pre-mRNA targets and *ASCO* lncRNA (Bardou *et al*, 2014). More recently, a RIP-seq approach on an NSRa fusion protein in *A. thaliana* mutant background allowed the identification of genome-wide NSR targets, e.g. specific alternatively spliced mRNAs as well as a plethora of lncRNAs, including *ASCO* (Bazin *et al*, 2018). Strikingly, *ASCO* was detected albeit not among the most abundant lncRNA, suggesting the existence of an intricate network of multiple lncRNAs and splicing factors interactions. In fact, we showed here that the impact of *ASCO* deregulation on AS at genome-wide level barely overlaps with the defects observed in the *nsra/b* mutant background (with or without auxin), indicating that *ASCO* and NSRs participate in common as well as in independent molecular mechanisms related to AS.

### The *ASCO* lncRNA knocked-down plants show altered sensitivity to flagellin

The comparison of the transcriptome of RNAi-*ASCO* and *35S:ASCO* plants reveals common and specific subsets of DAS genes. This dual effect caused by the up- or down-regulation of *ASCO* accumulation hints to the potential relevance of a stoichiometric factor impacting the action of *ASCO* within the spliceosome. *ASCO* silenced plants exhibit an enhanced sensitivity to flg22, in contrast to *35S:ASCO* and *nsra/b* plants. In agreement, overexpressing *ASCO* plants and *nsra/b* mutants behave similarly in response to auxin (Bardou *et al*, 2014). Our results suggest that *ASCO* has a wider function that the simple titration of NSR activity. Remarkably, we now determined that *ASCO* is recognized by additional splicing factors: the spliceosome core components PRP8a and SmD1b. Accordingly, RNAi-*ASCO* lines and a *prp8a* leaky mutant exhibit similar AS defects of flg22-regulated genes. However, *smd1b* mutants resulted in a deregulated ratio of isoforms of only *ESP* and *SR34*, but not *NUDT7*. The milder effect of *ASCO*-related SmD1b over the subset of pathogen-related genes may be due to a compensatory role of SmD1a in the *smd1b* background. Moreover, core splicing factors null mutants usually gave very severe phenotypes or embryo lethality and *smd1b* and *prp8a* mutants are leaky alleles with partial effects on global constitutive splicing. Altogether, our results indicate that a complex network of lncRNAs and splicing factors involving *ASCO*, PRP8a, SmD1b and NSRs dynamically shapes transcriptome diversity, integrating developmental and environmental cues, thus conditioning the response to biotic stress (Fig 7). In Arabidopsis, the lncRNA *ELF18-INDUCED LONG-NONCODING RNA 1* (*ELENA1*) is regulated by the perception of the translation elongation factor Tu (elf18) and it was identified as a factor enhancing resistance against *Pseudomonas syringae*. It was shown that *ELENA1* directly interacts with Mediator subunit 19a (MED19a), modulating the enrichment of MED19a on the *PATHOGENESIS-RELATED GENE1* (*PR1*) promoter (Seo *et al*, 2017). Several other examples of lncRNAs mediating the environmental control of gene expression illustrate the relevance of the noncoding transcriptome as a key integration factor between developmental and external cues (Marquadt et al., 2014; Heo and Sung 2011; Kim and Sung 2017; Kindgren et al 2018.) The sensitivity to pathogens has been shown to be affected in spliceosome-related mutants. For instance, it was recently reported that the *prp40c* mutants display an enhanced tolerance to *Pseudomonas syringae* (Hernando et al 2019, FPS). On the other hand, several other siplicing related genes have been identified as positive regulators of plant immunity against Pseudomonas (Jacqueline Monaghan et al., 2009 PLOS Pathogens; Palma, Zhao, Cheng, Bi, & Monaghan, 2007 Genes and Dev; F. Xu, Xu, Wiermer, Zhang, & Li, 2012 Plant Journal).

**Figure 7.**
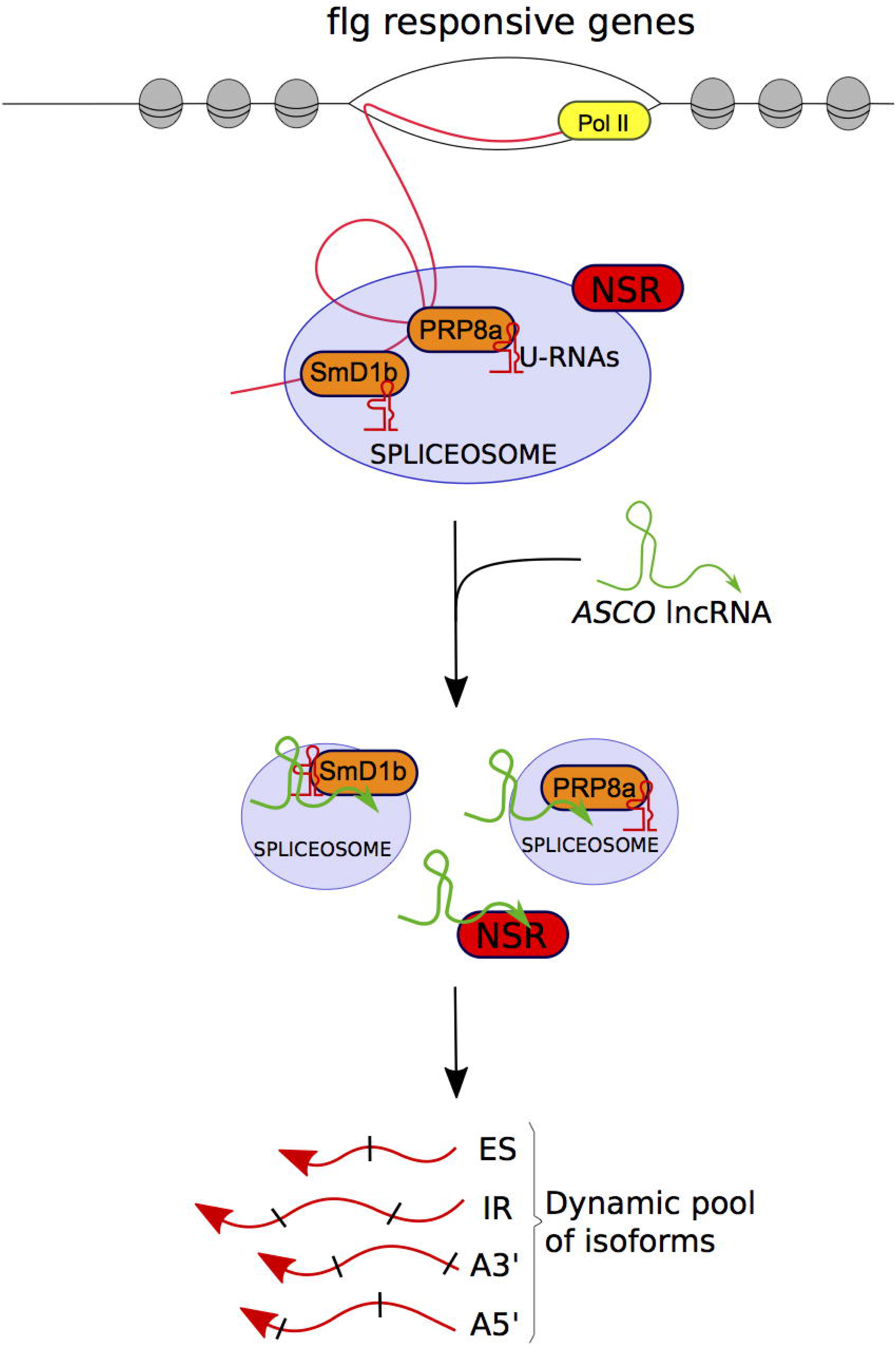
The interaction of *ASCO* lncRNA and the spliceosome components PRP8a and SmD1b shapes the transcriptional response to flg22 modulating alternative splicing. Proposed mechanism of *ASCO* lncRNA action. *ASCO* hijacks NSR proteins to modulate the population of alternatively spliced transcripts. Additionally, *ASCO* is recognized by PRP8a and SmD1b, two core components of the spliceosome, conditioning the SmD1b/PRP8a-dependent transcriptome diversity in response to flagellin.

### LncRNAs as highly variable components of the conserved spliceosomal machinery

Here we show that the highly structured lncRNA *ASCO*, which does not contain introns, is capable to interact with PRP8a and modulate PRP8a binding to *ASCO*-related AS targets. The spliceosome is a large complex composed of five different small nuclear ribonucleoprotein complexes subunits (snRNPs). Each subunit includes noncoding and non-polyadenylated small nuclear uridine (U)-rich RNAs (U snRNAs) and core spliceosomal proteins, along with more than 200 non-snRNPs splicing factors (Herold *et al*, 2009). PRP8a is one of the largest and most highly conserved proteins in the nucleus of eukaryotic organisms. It occupies a central position in the catalytic core of the spliceosome and has been implicated in several crucial molecular rearrangements (Grainger and Beggs, 2005). In *Arabidopsis*, analysis of *PRP8a* leaky mutation suggests that PRP8a recognizes the lncRNA *COOLAIR in vivo* to modulate its AS (Marquardt *et al*, 2014) hinting an interaction with lncRNAs. *COOLAIR* designates a set of transcripts expressed in antisense orientation of the locus encoding the floral repressor *FLC* (Whittaker and Dean, 2017). Two main classes of *COOLAIR* lncRNAs are produced by AS and polyadenylation of antisense transcripts generated from the *FLC* locus. One uses a proximal splice site and a polyadenylation site located in intron 6 of *FLC*, whereas the distal one results from the use of a distal splice and polyadenylation sites located in the *FLC* promoter (Whittaker and Dean, 2017). Notably, *prp8a* partial loss of function leads to a reduced usage of *COOLAIR* proximal polyadenylation site and an increase of *FLC* transcription which is associated with late flowering phenotypes (Marquardt *et al*, 2014). Interestingly, the *FLC/COOLAIR* module is strongly deregulated in the *nsra/b* double mutant and NSRa was linked to flowering time further supporting multiple interactions of lncRNAs and the splicing machinery (Bazin *et al*, 2018). Although NSRa-*COOLAIR* interaction seems not to occur, it was proposed that the control of NSRa over *COOLAIR* involves the direct interaction and processing of the polyadenylation regulatory gene *FPA* (Bazin *et al*, 2018). In the model legume *Medicago truncatula*, the NSRs closest homolog, RNA-BINDING PROTEIN 1 (RBP1), is localized in nuclear speckles where many components of the splicing machinery are hosted in plant cells. Remarkably, RBP1 interacts with a highly structured lncRNA, *EARLY NODULIN 40* (*ENOD40*), which participates in root symbiotic nodule organogenesis (Crespi *et al*, 1994; Charon *et al*, 1999; Campalans *et al*, 2004). *ENOD40* is highly conserved among legumes and was also found in other species such as rice (*Oryza sativa*; Gultyaev and Roussis, 2007), but shows no homology to *ASCO* lncRNA (Bardou *et al*, 2014). In contrast to the nuclear localization of *Arabidopsis ASCO*, *ENOD40* was found both in the nucleus and the cytoplasm, and it is able to relocalize RBP1 from nuclear speckles into cytoplasmic granules during nodulation. These observations hint a role of the lncRNA *ENOD40* in nucleocytoplasmic trafficking, potentially modulating RBP1-dependent splicing (Campalans *et al*, 2004) and further supporting the multiple interactions of lncRNAs with splicing regulators. A major result shown here is that *ASCO* is recognized by PRP8a and SmD1b, two central regulators of splicing and not only the NSR proteins which are plant-specific “peripheral” regulators of splicing. Indeed, the *nsra/b* null double mutants did not display major phenotypes in contrast to null PRP or SmD components. The identification of how the *ASCO* lncRNA interacts with PRP8a will certainly contribute to decipher the intricate network of lncRNAs fine-tuning the activity of core splicing factors, thus opening wide perspectives for the use of lncRNAs in the modulation of the dynamic population of alternatively spliced mRNAs in higher organisms. Interestingly, a search for *ASCO* homologues across the Brassicaceae family reveals that 9 additional copies of *ASCO* exist in *A. thaliana* and related sequences are also present in other Brassicaceae species, including *A. halleri*, *A. lyrata* and the more distant species *Capsella rubella* and *Capsella grandiflora* (Fig EV 11 A). However, none of the four detectable *A. thaliana ASCO*-like homologues suffered any significant alteration in RNAi-*ASCO* and *35S:ASCO* lines (Fig EV 11B), suggesting that none of them seem to compensate for the absence or presence of the *ASCO* lncRNA. The existence of *ASCO*-like sequences in other species suggest that conserved lncRNA-mediated mechanisms of AS regulation may occur through the interaction with highly conserved splicing factors. As PRP8a and SmD1b as well as the snRNAs are highly conserved spliceosomal components in contrast to the outstanding variability of lncRNA sequences along evolution, our results hint to a yet undiscovered evolutionary layer in the regulation of AS in different cell types and environmental conditions without affecting essential splicing activity. Structure and short sequences inside lncRNAs may contribute to the evolution of splicing regulatory networks in eukaryotes.

## Materials and Methods

### Plant material and growth conditions

All the lines used in this study were in the *A. thaliana* Columbia-0 (Col-0) background. We used the *nsra/b* double mutant and the *ASCO* overexpressing lines from Bardou *et al* (2014). The insertion lines WiscDsLoxHs110_08A (*asco-1)* and SAIL_812_C08 (*asco-2)* were obtained from the T-DNA mutant collection at the Salk Institute Genomics Analysis Laboratory (SIGnAL, http://signal.salk.edu/cgi-bin/tdnaexpress) via NASC (http://arabidopsis.info/). Seeds from *prp8-7* (Sasaki *et al*, 2015) and *smd1b* (Elvira-Matelot *et al*, 2016) mutants were provided by H. Vaucheret. The *pUBQ10:SmD1b-GFP* line (Elvira-Matelot *et al*, 2016) was used for RNA immunoprecipitation assays. Plants were grown at 20°C with a 16-h light/ 8-h dark photoperiod (long days) on solid half-strength Murashige and Skoog (1/2MS) medium.

### Generation of transgenic lines

#### Pro*ASCO::GUS transgenic lines*

The promoter region of *ASCO* (2631-bp upstream of the transcription start) was amplified from *A. thaliana* genomic DNA using gene-specific primers listed in Table EV5. The amplicon was subcloned into the pENTR/D-TOPO vector and recombined in a pKGWFS7 binary destination vector, upstream of the *GFP* and GUS sequences. Pro*ASCO*::*GUS* constructs were transferred into *A. thaliana* by standard Agrobacterium-mediated protocol (Clough and Bent, 1998). Three lines were selected based on 3:1 segregation for the transgene (single insertion) and brought to T3 generation where the transgene was in a homozygous state. All lines behave similarly as for *GUS* expression.

#### RNAi*-ASCO knocked-down lines*

The first 233-bp of the *ASCO* transcript was amplified from *A. thaliana* genomic DNA using gene-specific primers listed in Table EV5. Amplicons were sub-cloned into the pENTR/D-TOPO vector and recombined in a pFRN binary destination vector (Ariel *et al*, 2012) to target the *ASCO* RNA by long dsRNA hairpin formation.

### Root growth analysis

For analysis of auxin impact on root architecture, plants were grown as described in Bardou *et al* (2014). Briefly, seeds were sterilized and directly sown on plates containing 1/2MS medium supplemented or not with 100 nM NAA. Plantlet root architecture was analyzed using the RootNav software after 7 days of growth (Pound *et al*, 2013). For analysis of flagellin impact on root architecture, plants were previously grown 5 days on solid 1/2MS medium + 1% sucrose and then transferred for additional 9 days in liquid 1/2MS media + 1% sucrose supplemented or not with 0.1 μΜ or 1 μΜ of synthetic flg22 peptide (GeneCust). For each plantlet, lateral roots were counted, and the primary root length was measured using the RootNav software. Experiments were done at least 2 times, with a minimum of 16 plants per genotype and condition. Statistical tests were performed using the Mann Whitney’s U test (p< 0.05).

### Root meristem measurements

Plants were grown as for root growth analysis in response to flg22. The treated plants were stained with SCRI Renaissance 2200 (Renaissance Chemicals) as described in Musielak *et al* (2015). Images were obtained with LSM880 (Zeiss) confocal microscope. The SR2200 fluorescence was excited with a 405 nm laser line and emission recorded between 410 and 686 nm (405/410-686). Cell counting and primary root meristem measurements were performed using ImageJ package (https://imagej.nih.gov/ij/). Experiments were done 2 times, with a minimum of 18 plants per genotype and condition. Statistical tests were performed using the Student’s T test (p< 0.05).

### Histochemical GUS staining

Histochemical GUS staining was performed according to Latrasse *et al* (2013). Briefly, 10-day-old plantlets grown in standard conditions were fixed in cold 90% acetone and incubated overnight at 37°C in the GUS staining buffer. Roots were subsequently fixed in 4% paraformaldehyde for 1 h and washed several times in 70% ethanol before a final wash in 10% glycerol prior observation. Images were acquired using an AxioImagerZ2 microscope (Zeiss).

### RNA extraction and RT-PCR analyses

Total RNA was extracted using Quick-RNA Kit (ZYMO RESEARCH) and DNase treatment was performed according to the manufacturer’s protocol. One μg of DNase-free RNA was reverse transcribed using Maxima H Minus Reverse Transcriptase (Thermo Scientific). cDNA was then amplified in RT-qPCRs using LightCycler 480 SYBR Green I Master (Roche) and transcript-specific primers on a Roche LightCycler 480 thermocycler following standard protocol (45 cycles, 60°C annealing). Experiments were done in biological triplicates with at least three technical replicates. Expression was normalized to 2 constitutive genes (AT1G13320 and AT4G26410; Wang *et al*, 2014). For analysis of flg22 impact on gene expression, plants were previously grown 9 days on solid 1/2MS medium + 1% sucrose and then transferred for additional 24 h in liquid 1/2MS medium + 1% sucrose before adding or not 1 μΜ of flg22. Five plantlets were pooled for each replicate. The fold induction of expression after flg22 treatment was normalized to the WT response considered as 100%. For analysis of gene expression after a flg22 kinetic, roots from 8 plants were pooled for each replicate. For AS analysis, isoform specific primers were designed for each differential event and the signal was normalized with respect to an internal gene probe (called INPUT) corresponding to a common exon for each group of transcripts. This allows differentiating the change of each isoforms independently of the expression level of the studied gene (splicing index) in each sample (Tran *et al*, 2016). The splicing index was calculated following this equation : splicing index = 2[^ct^(^s^P^ecific isoform^) -(ACt(iNPUT)]. Error bars on qRT-PCR experiments represent standard deviations, and significant differences were determined using a student’s T test (p <= 0.05, n >= 3) (n = 3). All the used primers are listed in Table EV5.

For RT-PCR analysis, the amplification was performed using Phusion High-Fidelity DNA Polymerase and transcript-specific primers as manufacturer’s protocol. PCR products were separated on an 8% polyacrylamide gel stained with SYBR Gold (Thermo Fischer Scientific) and revealed using a ChemiDoc MP Imaging System (Biorad). Band intensity was quantified using ImageJ package (https://imagej.nih.gov/ij/). Isoform ratio was calculated as the ratio of intensity of the two bands corresponding either to the alternatively-spliced or spliced transcript isoforms, respectively.

### Transcriptome studies

Total RNA was extracted using RNeasy plant mini kit (Qiagen) from whole 14-day-old Col-0, *35S:ASCO1* and RNAi-*ASCO1* plants grown on 1/2MS medium. Three independent biological replicates were produced per genotype. For each biological repetition and each point, RNA samples were obtained by pooling RNA from more than 200 plants. After RNA extraction, polyA RNAs were purified using Dynabeads mRNA direct micro kit (Ambion). Libraries were constructed using the Tru-Seq Stranded mRNA Sample Prep kit (Illumina®). Sequencing was carried out at the POPS Transcriptomic Platform, Institute of Plant Sciences Paris-Saclay in Orsay, France. The Illumina HiSeq2000 technology was used to perform paired-end 100-bp sequencing. A minimum of 30 million of paired-end reads by sample were generated. RNA-seq preprocessing included trimming library adapters and quality controls with Trimmomatic (Bolger *et al*, 2014). Paired-end reads with Phred Quality Score Qscore > 20 and read length > 30 bases were kept, and ribosomal RNA sequences were removed with SortMeRNA (Kopylova *et al*, 2012). Processed reads were aligned using Tophat2 with the following arguments : --max-multihits 1 -i 20 --min-segment-intron 20 --min-coverage-intron 20 --library-type fr-firststrand --microexon-search -I 1000 --max-segment-intron 1000 --max-coverage-intron 1000 -b2-very- sensitive. Reads overlappings exons per genes were counted using the FeatureCounts function of the Rsubreads package using the GTF annotation files from the Araport11 repository (https://www.araport.org/downloads/Araport11_Release_201606/annotation/Araport11_GFF3_genes_transposons.201606.gff.gz). Significance of differential gene expression was estimated using DEseq2 (Love et al 2019), and the FDR correction of the p-value was used during pairwise comparison between genotypes. A gene was declared differentially expressed if its adjusted p-value (FDR) was ≤ 0.01 and its absolute fold change was ≥ 1.5.

### Gene Ontology analysis

Gene ontology enrichment analysis was done using AgriGO (http://bioinfo.cau.edu.cn/agriGO) and default parameters.

### AS analysis

The RNA profile analysis was performed using the RNAprof software (v1.2.6) according to Tran *et al* (2016). Briefly, RNAprof software allows detection of differential RNA processing events from the comparison of nucleotide level RNA-seq coverage normalized for change in gene expression between conditions. Here, the RNAprof analysis compared RNA-seq data from biological triplicates of WT, RNAi-*ASCO1* and *35S:ASCO1* lines. Differentially processed regions genes were filtered as follows: fold change > 2; p-value < 0.001. Overlap between gene features and differentially processed regions was done using in-house R scripts (https://github.com/JBazinIPS2/Bioinfo/blob/master/RNAprof_events_selection.Rrst). Only regions fully included in a gene features were kept for further analysis.The RNAprof software archive, including documentation and test sets, is available at the following address: http://rna.igmors.u-psud.fr/Software/rnaprof.php. Transcript level quantification was performed using pseudo-alignment counts with kallisto (Bray *et al*, 2016) on AtRTD2 transcripts sequences (https://ics.hutton.ac.uk/atRTD/RTD2/AtRTDv2_QUASI_19April2016.fa) with a K-mer size of 31-nt. Differential AS events in the AtRTD2 database were detected using SUPPA2 (Trincado *et al*, 2018) with default parameters. Only events with an adjusted p-val < 0.01 were kept for further analysis. Isoforms switch identification was performed with the IsoformSwitchAnalyzeR package (Vitting-Seerup and Sandelin 2019) according to Bazin *et al* (2018).

### Whole-mount immunolocalization

Specific rabbit polyclonal antibodies were developed against PRP8a using the peptide TNKEKRERKVYDDED (Li International). Five-day-old seedlings were fixed with 4% paraformaldehyde in MicroTubule Stabilization Buffer (MTSB; Pasternak *et al*, 2015) for 1 h and rinsed once in glycine 0.1 M and twice with MTSB. Cell walls were partially digested for 45 min at 37°C in cellulase R10 1% w/v (Onozuka), pectolyase 1% w/v and cytohelicase 0.5% w/v (Sigma) solution. After two PBS washes, root tissues were squashed on polylysine-treated glass slides (VWR International) and dipped in liquid nitrogen. The coverslip was then removed, and the slides were left to dry. After 2 rinses with PBS and 2 with PBS-0.1% triton, they were treated with BSA 3% in PBS-Triton buffer for 1 h and incubated with the anti-PRP8a antibody (dilution 1:400) for 16 h at 4°C in a humid chamber. After incubation, slides were rinsed 8-10 times with PBS-Triton and incubated for 1 h at 37°C with the secondary antibody (anti Rabbit IgG coupled to Alexa Fluor® 594, dilution 1:500; Thermo Scientific) and rinsed 10 times with PBS-Triton and once with PBS. Slides were mounted in Vectashield© containing DAPI (VECTOR Laboratories). Images were obtained with LSM880 (Zeiss) confocal microscope equipped with Plan-Apochromat 63x/NA1,40 Oil M27 lens. Dapi and Alexa 594 fluorescences were respectively excited with 405 nm and 561 nm diodes and recorded between 410-500 nm and 570-695 nm.

### LncRNA-bound nuclear protein isolation by RNA purification

A method adapted from the ChIRP protocol (Chu *et al*, 2011; Ariel *et al*, 2014; Chu and Chang, 2016) was developed to allow identification of nuclear proteins bound to specific lncRNAs. Briefly, plants were *in vivo* crosslinked, and nuclei of cells purified and extracted through sonication. The resulting supernatant was hybridized against biotinylated complementary oligonucleotides that tile the lncRNA of interest and putative lncRNA-containing protein complexes were isolated using magnetic streptavidin beads. Co-purified ribonucleoprotein complexes were eluted and used to purify RNA or proteins, which were later subject to downstream assays for identification and quantification.

#### Probe design

Antisense 20-nt oligonucleotide probes were designed against the *ASCO* full-length sequence (AT1G67105) using an online designer at http://singlemoleculefish.com/. All probes were compared with the *A. thaliana* genome using the BLAST tool at the NCBI, and probes returning noticeable homology to non-*ASCO* targets were discarded. Eighteen probes were finally generated and split into two sets based on their relative positions along the *ASCO* sequence, such as EVEN-numbered and ODD-numbered probes were separately pooled. A symmetrical set of probes against *LacZ* RNA (Ariel *et al*, 2014) was also used as the mock control. All probes were ordered biotinylated at the 3’ end (Invitrogen).

#### Crosslinking and ribonucleoprotein complexes purification

For protein extraction, approximately 250 g of 7-day-old Col-0 plants grown on solid half-strength MS medium were irradiated three times with UV using a CROSSLINKER® CL-508 (Uvitec) at 0.400 J/cm^2^. For RNA extraction, 10 g of 7-day-old Col-0 plants grown on solid half-strength MS medium were crosslinked under vacuum for 15 min with 37 mL of 1% (v/v) formaldehyde. The reaction was stopped by adding 2.5 mL of 2 M glycine and seedlings were rinsed with Milli-Q purified water. For both crosslinking methods, 6g of the fixed material was ground in liquid nitrogen and added to 50 mL tubes with 25 mL of Extraction Buffer 1 to 15 ml of plant material ground to fine dust (the nuclei were prepared starting with 30 50ml-tube; Buffer 1: 10 mM Tris-HCl pH 8, 0.4 M sucrose, 10 mM MgCl_2_, 5 mM β-mercaptoethanol, 1 mL/30 g of sample powder Protease Inhibitor Sigma Plant P9599). The solution was then filtered through Miracloth membrane (Sefar) into a new tube and 5 mL of Extraction Buffer 2 (10 mM Tris-HCl pH 8, 0.25 M sucrose, 10 mM MgCl_2_, 5 mM β-mercaptoethanol, 1% Triton X-100, 50 μL Protease Inhibitor) were added. The solution was then centrifuged, the supernatant discarded and 500 μL of Extraction Buffer 3 (10 mM Tris-HCl pH 8, 1.7 M sucrose, 2 mM MgCl_2_, 5 mM β-mercaptoethanol, 0.15% Triton X-100, 50 μL Protease Inhibitor) and layered on top of fresh Extraction Buffer 3 in a new tube. After centrifugation at 13000 rpm for 2 min at 4°C to pellet nuclei, the supernatant was discarded and the pellet resuspended in 300 μL of Nuclei Lysis Buffer (50 mM Tris-HCl pH 7, 1% SDS, 10 mM EDT,; 1 mM DTT, 50 μL Protease Inhibitor, 10 μL RNAse Inhibitor per tube) to degrade nuclear membranes. Samples were sonicated three times in refrigerated BIORUPTOR Plus (Diagenode), 10 cycles 30 sec ON - 30 sec OFF in a Diagenode TPX microtube M-50001. After centrifugation, the supernatant was transferred to a new tube and diluted two times volume in Hybridization Buffer (50 mM Tris-HCl pH 7, 750 mM NaCl, 1% SDS, 15% Formamide, 1 mM DTT, 50 μL Protease Inhibitor, 10 μL RNAse Inhibitor). One hundred pmol of probes were added to samples and incubated 4 h at 50°C in a thermocycler. Samples were transferred to tubes containing Dynabeads-Streptavidin C1 (Thermo Fisher Scientific) and incubated 1 h at 50°C. Then samples were placed on a magnetic field and washed three times with 1 mL of Wash Buffer (2x SSC, 0.5% SDS, 1 mM DTT, 100 μL Protease Inhibitor).

#### Protein purification

Samples for protein extraction were DNase-treated according to the manufacturer (Thermo Scientific). After addition of 1.8 mL of TCA-Acetone (5 mL 6.1 N TCA + 45 mL Acetone + 35 μL β-mercaptoethanol), samples were incubated overnight at −80°C. After centrifugation at 20000 rpm for 20 min and 4°C, the supernatant was discarded and 1.8 mL of Acetone Wash Buffer (120 mL Acetone, 84 pL β-mercaptoethanol) was added to the samples. Then, samples were incubated 1 h at −20°C, and centrifuged again at 20000 rpm for 20 min and 4°C. The supernatant was discarded, and the dry pellet was used for Mass Spectrometry analysis.

#### RNA purification

Samples for RNA extraction were boiled for 15 min after washing with 1 mL of Wash Buffer (2X SSC, 0.5% SDS, 1 mM DTT, 100 μL Protease Inhibitor). Beads were removed in a magnetic field and Trizol/Chloroform RNA extraction was performed according to the manufacturer (Sigma). RNAs were precipitated using 2 volumes EtOH 100%, 10% 3 M sodium acetate and 1 pL glycogen, and washed with EtOH 70%. RNAs were kept at −20°C before use for reverse transcription and RT-qPCR analysis.

### Liquid Chromatography - Mass Spectrometry (LC-MS) analysis

Proteins purified from ribonucleoprotein complexes were analyzed using the King Abdullah University of Science and Technology proteomic facilities. Dry pellets of samples purified with either ODD, EVEN or *LacZ* probe sets were solubilized in Trypsin Buffer (Promega) for digestion into small peptides. The solubilized peptides were then injected into a Q Exactive™ HF hybrid quadrupole-Orbitrap mass spectrometer (Thermo Scientific) with a Liquid Chromatography (LC) Acclaim PepMap C18 column (25 cm length x 75 μm I.D. x 3 μm particle size, 100 Â porosity, Dionex). Data were analyzed for each sample using Mascot software (Matrix Science), with a minimal sensitivity of 2 detected peptides per identified protein.

### RNA immunoprecipitation

Eleven-day-old plants grown in Petri dishes were irradiated three times with UV using a CL-508 cross-linker (Uvitec) at 0.400 J/cm^2^. Briefly, fixed material was ground in liquid nitrogen and homogenized and nuclei isolated and lysed according to Gendrel *et al* (2005). RNA immunoprecipitation was basically performed as described by Carlotto *et al* (2015). The nuclei extract (input) was used for immunoprecipitation with 50 μL of Dynabeads-Protein A (Thermo Fisher Scientific) and 1 μg of anti-GFP antibodies (Abcam ab290) or anti-PRP8a (Li International), respectively. Beads were washed twice for 5 min at 4°C with wash buffer 1 (150 mM NaCl, 1% Triton X-100, 0.5% Nonidet P-40, 1 mM EDTA, and 20 mM Tris-HCl, pH 7.5) and twice with wash buffer 2 (20 mM Tris-HCl, pH 8) and finally resuspended in 100 μL Proteinase K buffer (100 mM Tris-HCl, pH 7.4, 50 mM NaCl, and 10 mM EDTA). Twenty microliters were saved for further immunoblot analysis. After Proteinase K (Ambion AM2546) treatment, beads were removed with a magnet, and the supernatants were transferred to a 2 mL tube. RNA was extracted using TriReagent (Sigma-Aldrich) as indicated by the manufacturer. Eighty microliters of nuclei extracts was used for input RNA extraction. The immunoprecipitation and input samples were treated with DNase, and random hexamers were used for subsequent RT. Quantitative real-time PCR reactions were performed using specific primers (Table EV5). Results were expressed as a percentage of cDNA detected after immunoprecipitation, taking the input sample as 100%. For immunoblot analysis, immunoprecipitated proteins were eluted by incubating 20 μL of beads in 20 μL of 2x SDS-loading buffer without β-mercaptoethanol (100 mM Tris-HCl pH 6.8, 20% glycerol, 12.5 mM EDTA, 0.02% bromophenol blue) at 50°C for 15 min. Input, unbound and eluted fractions were boiled in 2x SDS loading buffer with 1% β-mercaptoethanol for 10 min, loaded onto a 4-20% Mini-PROTEAN® TGX™ Precast Protein Gels (Biorad) and transferred onto a PVDF membrane using the Mini Trans-Blot® Cell system (BioRad) for 3 h at 70 V. Membranes were blocked in 5% dry non-fat milk in PBST and probed using PRP8a antibody (1:500) and an HRP coupled anti-Rabbit secondary antibody (Biorad, 1:10000). All antibodies were diluted in 1% dry nonfat milk in PBST. Blots were revealed with the Clarity ECL substrate according the manufacturer instruction (BioRad) and imaged using the ChemiDoc system (BioRad).

### Data deposition

Data was deposited in CATdb database (Gagnot *et al*, 2008; http://tools.ips2.u-psud.fr/CATdb/) with ProjectID NGS2016-07-ASCOncRNA. This project was submitted from CATdb into the international repository GEO (http://www.ncbi.nlm.nih.gov/geo) with ProjetID GSE135376.

## Acknowledgements

Synthetic flg22 peptide was kindly provided by J. Colcombet (IPS2). Seeds from the *prp8-7* (Sasaki *et al*, 2015) and the *smdlb* (Elvira-Matelot *et al*, 2016) leaky mutants were kindly provided by H. Vaucheret. This work was supported by grants from ANPCyT (PICT 2016-0289 and -0007, Argentina), CNRS (Laboratoire International Associé NOCOSYM), the ‘Laboratoire d’Excellence (LABEX)’ Saclay Plant Sciences (SPS; ANR-10-LABX-40), the Ministère Français de l’Enseignement Supérieur et de la Recherche (RR) and the ANR grants (ANR-15-CE20-0002-01 EPISYM and ANR 16-CE20-0003-04 SPLISIL) as well as the EPIMMUNITY International project between IPS2, France and KAUST University, Saudi Arabia. The POPS platform benefits from the support of the LabEx Saclay Plant Sciences-SPS (ANR-10-LABX-0040-SPS).

## Author contributions

RR, JB, NRB, MM, LL, AC, CC and FA performed the experiments; RR, JB, CC, MB, MC and FA analyzed the data; JB performed bioinformatic analyses; JB, MC and FA designed the experiments; MC and FA conceived the study and wrote the manuscript.

## Expanded View Figure Legends

**Figure EV1. The response to flg22 is altered in RNAi-*ASCO* lines.**

**A** *ASCO* transcript levels in 2 independent 14-day-old RNAi-*ASCO* lines compared to WT.

**B** RNA-seq read coverage on *ASCO* lncRNA in RNAi-*ASCO* and WT seedlings RNA-seq data. The region cloned to generate dsRNA is indicated on the gene structure. Coverage plots were made with IGV software (Robinson et al., 2018) using normalized (read per million) bigwig files. The same scale (0-200) was used for each track.

**C, D** Primary root length (**C**) and lateral root density (**D**) of WT and two independent RNAi-*ASCO* lines grown on media with or without 100 nM NAA and measured 7 days after germination. The asterisk (*) indicates a significant difference as determined by Mann Whitney’s U (test < 0.05, n = 16).

**E** Scheme of the transcriptional fusion to *GFP:GUS* used to analyze the *ASCO* promoter activity.

**F** Activity of *proASCO* during LR development. Scale bar corresponds to 20 μm.

**Figure EV2 Expression of immunity-related TF in RNAi-ASCO lines and ASCO expression in roots during flg22 treatment.**

**A** Transcript levels of stress-related DEG identified in the RNA-seq data in the WT and the two independent RNAi-*ASCO* lines, measured by RT-qPCR. RNAs were extracted from 14-day-old plants. Data are mean of 3 independent biological replicates. Error bars represent standard deviation. The asterisk (*) indicates a significant difference as determined by Student’s T (test < 0.05, n = 3).

**B** Time-course analysis of *ASCO* and *CYP81F2* (AT5G57220) expression levels after treatment with media supplemented or not with 1 μΜ flg22. RNAs were extracted from 10-day-old plants. Data are mean of 3 independent biological replicates. Error bars represent standard deviation.

**Figure EV3 The response to flg22 is altered in RNAi-*ASCO* lines.**

**A** A representative picture of 14-day-old plants grown 9 days in liquid 1/2MS corresponding to Mock condition.

**B** A representative picture of root apical meristems after cell wall staining, in Mock condition. TZ: Transition Zone; QC: Quiescent Center.

**C** Primary root length of WT and two independent RNAi-*ASCO* lines 9 days after transfer in 1/2MS supplemented with 0.1 μΜ or 1 μΜ flg22.

**D** Mean of meristematic cell number in WT and RNAi-*ASCO*1 primary root apex, treated or not with 1 μΜ flg22. In (**C**) and (**D**) error bars indicate the standard error. The asterisk (*) indicates a significant difference as determined by Mann Whitney’s U test (p < 0.05, n = 18).

**E** Transcript levels of a subset of flg22 responsive genes (AGIs are indicated in Table EV5) in control conditions.

**F** Relative modulation of the same genes as in **D**, in response to flg22. Results are normalized against the response in WT plants, taken as 100%. The asterisk (*) indicates a significant difference as determined by Student’s T test (p < 0.05, n = 3).

**Figure EV4 *ASCO* insertion mutants do not exhibit strong expression changes**

**A** Scheme of T-DNA insertions along *ASCO* locus in *asco-1* and *asco-2* mutants. Red arrows indicate probes used for qPCR expression analysis.

**B** *ASCO* transcript levels in 2 independent 14-day-old 35-*ASCO* lines, *asco-1* and *asco-2* compared to WT.

**C** Quantification of *ASCO* transcript levels targeting different regions of the locus by RT-qPCR in WT and *asco* mutants. A, B, C and D probe positions are indicated in **A**.

**D, E** Primary root length (**D**) and lateral root density (**E**) of WT, *asco-1* and *asco-2* plants 9 days after transfer in 1/2MS supplemented with 1 μΜ flg22.

**Figure EV5 Differential RNA processing events identified in RNAi-*ASCO*.**

**A, B, C, D, E, F** Differential RNA processing events of *SR34* (AT1G02840, **A**), *NUDT7* (AT4G12720, **B**), *NRG* (AT2G29290, **C**), *SEN1* (AT4G35770, **D**), *SNC4* (AT1G66980, **E**) and *RPP4* (AT4G16860, **F**) transcripts detected by RNAprof from the comparison of RNA-seq libraries of 14-day-old plants. Comparative transcriptional abundance are indicated for WT (red) and RNAi-*ASCO*1 (blue) plants. Three biological replicates were used for the analyses. Vertical purple lines and p-values indicate significant differential processing events. Each profile is associated with the structure of the corresponding RNA isoforms. Large black boxes indicate exons, narrow black boxes indicate UTRs and black lines indicate introns.

**Figure EV6. Validation of AS events in RNAi-*ASCO* lines.**

**A, D, G, J, M** Differential RNA processing events of *ESP* (AT1G54040), *NRG, SEN1, SNC4* and *RPP4* transcripts detected by IsoSwitch (*ESP*) or RNAprof *(NRG, SEN1, SNC4 and RPP4)* from the comparison of mRNA-seq libraries of 14-day-old plants. For *NRG, SEN1, SNC4 and RPP4,* comparative transcriptional abundance are indicated for WT (red) and RNAi-*ASCO*1 (blue) plants. Three biological replicates were used for the analyses. Vertical purple lines in **D**, **G, J** and **M** and p-values indicate significant differential processing events. The structure of each associated RNA isoform is represented as follows : large black boxes indicate exons, narrow black boxes indicate UTRs, black lines indicate introns. Red arrows indicate probes used for gel electrophoresis. Colored boxes indicate protein domains affected by an AS event.

**B, E, H, K, N** Analyses of RT-PCR products of corresponding transcripts on 8% acrylamide gel.

**C, F, I, L, O** Quantification of the ratio of the corresponding isoforms detected in gels in **B, E, H, K** and **N**, respectively.

RNAs were extracted from WT and RNAi-*ASCO* 14-day-old plants. The asterisk (*) indicates a significant difference as determined by Student’s T test (p < 0.05, n = 3).

**Figure EV7. qPCR analysis of several AS events identified in RNAi-ASCO lines.**

**A, B, C** and **D** Quantification of *SR34* (**A**), *ESP* (**B**), *NUDT7* (**C**) and *SNC4* (**D**) isoforms by RT-qPCR. The asterisk (*) indicates a significant difference as determined by Student’s T test (p< 0.05, n = 3).

**Figure EV8 nsra/b affect root responses to flg22 and splicing of subsets of genes partially overlapping with RNAi-ASCO and 35S:ASCO**

**A** Overlap between differentially spliced events in RNAi-*ASCO*, *35S:ASCO* and WT and *nsra/b* mutant and WT treated with NAA (Bardou *et al*, 2014). **B,**Primary root length and **C**, lateral root density of WT and *nsra/b* double mutants 9 days after transfer in 1/2MS supplemented with 1 μΜ flg22. The asterisk (*) indicates a significant difference as determined by Mann Whitney’s U test (p < 0.05, n > 16)

**Figure EV9. Novel specific antibody against AtPRP8a confirmed the protein localization in the cell nucleus.**

**A** Whole-mount immunolocalization using the specific antibody against PRP8a. The signals of DAPI (grey channel) and AtPRP8a protein (red channel) colocalized in the nuclei of *A. thaliana* WT plants. The negative control corresponds to the immunolocalization assay without anti-AtPRP8a antibody. Scale bar represents 10 μm.

**Figure EV10. Analyses of *NUDT7* and *SNC4* in the *smd1b* background.**

**A, D**, **G** Structure of *NUDT7* (**A**, **G**) and *SNC4* (**D**) RNA isoforms. Large black boxes indicate exons, narrow black boxes indicate UTRs, black lines indicate introns. Red arrows indicate probes used for gel electrophoresis. Colored boxes indicate protein domains affected by an AS event.

**B, E, H** Analyses of RT-PCR products of corresponding transcripts on 8% acrylamide gel.

**C, F, I** Quantification of the ratio of the corresponding isoforms detected in gels in **B, E** and **H,** respectively. **J.** *SNC4* transcript levels in *smd1b* mutant compared to WT. RNAs were extracted from WT, *prp8a* and *smd1b* 14-day-old plants. The asterisk (*) indicates a significant difference as determined by Student’s T test (p < 0.05, n = 3).

**Figure EV11 Identification of ASCO homologs in A.thaliana and the Arabidopsis Brassicaceae family**

A Maximum likelihood tree depicting the evolutionary history of *ASCO* in the Brassicaceae family. Bootstrap support values are indicated above branches. Sequences identifiers are colored by species.

B Normalized expression level of detectable *ASCO*-like transcripts in WT (blue bars), RNAi-*ASCO* (green bars) and *35S:ASCO* (red bars) RNA-seq data.

## Appendix

**Table EV1. List of differentially expressed genes in WT vs RNAi-ASCO1**

**Table EV2. List of differential RNA processing events determined by RNAprof.**

**Table EV3. List of differential AS events determined by SUPPA.**

**Table EV4. List of differential RNA processing events determined by IsoformSwitchAnalyzeR package.**

**Table EV5. List of oligonucleotides used for this work.**

## References

Alamancos GP, Pages A, Trincado JL, Bellora N & Eyras E (2015) Leveraging transcript quantification for fast computation of alternative splicing profiles. RNA 21: 1521–1531

Ariel F, Brault-Hernandez M, Laffont C, Huault E, Brault M, Plet J, Moison M, Blanchet S, Ichante JL, Chabaud M, Carrere S, Crespi M, Chan RL & Frugier F (2012) Two direct targets of cytokinin signaling regulate symbiotic nodulation in Medicago truncatula. Plant Cell 24: 3838–3852

Ariel F, Jegu T, Latrasse D, Romero-Barrios N, Christ A, Benhamed M & Crespi M (2014) Noncoding transcription by alternative RNA polymerases dynamically regulates an auxin-driven chromatin loop. Mol. Cell 55: 383–396

Ariel F, Romero-Barrios N, Jegu T, Benhamed M & Crespi M (2015) Battles and hijacks: noncoding transcription in plants. Trends Plant Sci. 20: 362–371

Asai T, Tena G, Plotnikova J, Willmann MR, Chiu W-L, Gomez-Gomez L, Boller T, Ausubel FM & Sheen J (2002) MAP kinase signalling cascade in Arabidopsis innate immunity. Nature 415: 977–983

Bardou F, Ariel F, Simpson CG, Romero-Barrios N, Laporte P, Balzergue S, Brown JWS & Crespi M (2014) Long Noncoding RNA Modulates Alternative Splicing Regulators in Arabidopsis. Dev. Cell 30: 166–176

Barry G, Briggs JA, Vanichkina DP, Poth EM, Beveridge NJ, Ratnu VS, Nayler SP, Nones K, Hu J, Bredy TW, Nakagawa S, Rigo F, Taft RJ, Cairns MJ, Blackshaw S, Wolvetang EJ & Mattick JS (2014) The long non-coding RNA Gomafu is acutely regulated in response to neuronal activation and involved in schizophrenia-associated alternative splicing. Mol. Psychiatry 19: 486–494

Bazin J, Romero N, Rigo R, Charon C, Blein T, Ariel F & Crespi M (2018) Nuclear Speckle RNA Binding Proteins Remodel Alternative Splicing and the Non-coding Arabidopsis Transcriptome to Regulate a Crosstalk Between Auxin and Immune Responses. Front. Plant Sci. 9: 1209

Beck M, Wyrsch I, Strutt J, Wimalasekera R, Webb A, Boller T & Robatzek S (2014) Expression patterns of flagellin sensing 2 map to bacterial entry sites in plant shoots and roots. J. Exp. Bot. 65: 6487–6498

Beltran M, Puig I, Pena C, Garcia JM, Alvarez AB, Pena R, Bonilla F & de Herreros AG (2008) A natural antisense transcript regulates Zeb2/Sip1 gene expression during Snaill-induced epithelial-mesenchymal transition. Genes Dev. 22: 756–769

Bethke G, Unthan T, Uhrig JF, Poschl Y, Gust AA, Scheel D & Lee J (2009) Flg22 regulates the release of an ethylene response factor substrate from MAP kinase 6 in Arabidopsis thaliana via ethylene signaling. Proc. Natl. Acad. Sci. U. S. A. 106: 8067­8072

Bolger AM, Lohse M & Usadel B (2014) Trimmomatic: a flexible trimmer for Illumina sequence data. Bioinformatics 30: 2114–2120

Boon K-L, Grainger RJ, Ehsani P, Barrass JD, Auchynnikava T, Inglehearn CF & Beggs JD (2007) prp8 mutations that cause human retinitis pigmentosa lead to a U5 snRNP maturation defect in yeast. Nat. Struct. Mol. Biol. 14: 1077–1083

Boudsocq M, Willmann MR, McCormack M, Lee H, Shan L, He P, Bush J, Cheng S-H & Sheen J (2010) Differential innate immune signalling via Ca(2+) sensor protein kinases. Nature 464: 418–422

Bray NL, Pimentel H, Melsted P & Pachter L (2016) Near-optimal probabilistic RNA-seq quantification. Nat. Biotechnol. 34: 525–527.

Campalans A, Kondorosi A & Crespi M (2004) Enod40, a Short Open Reading Frame - Containing mRNA, Induces Cytoplasmic Localization of a Nuclear RNA Binding Protein in Medicago truncatula. Plant Cell 16: 1047–1059

Carlotto N, Wirth S, Furman N, Ferreira Solari N, Ariel F, Crespi M & Kobayashi K (2015) The chloroplastic DEVHDbox RNA helicase INCREASED SIZE EXCLUSION LIMIT 2 involved in plasmodesmata regulation is required for group II intron splicing. Plant Cell & Environment 39: 165–173

Charon C, Sousa C, Crespi M & Kondorosi A (1999) Alteration of enod40 Expression Modifies Medicago truncatula Root Nodule Development Induced by Sinorhizobium meliloti. Plant Cell 11: 1953–1965

Chaudhary S, Jabre I, Reddy ASN, Staiger D & Syed NH (2019) Perspective on Alternative Splicing and Proteome Complexity in Plants. Trends Plant Sci. 24: 496–506

Chu C, Qu K, Zhong FL, Artandi SE & Chang HY (2011) Genomic maps of long noncoding RNA occupancy reveal principles of RNA-chromatin interactions. Mol. Cell 44: 667­678

Chu C & Chang HY (2016) Understanding RNA-chromatin interactions using chromatin isolation by RNA purification (ChIRP). Methods Mol. Biol. 1480: 115–123

Claudius A-K, Romani P, Lamkemeyer T, Jindra M & Uhlirova M (2014) Unexpected role of the steroid-deficiency protein ecdysoneless in pre-mRNA splicing. PLoS Genet. 10: e1004287

Clough SJ & Bent AF (1998) Floral dip: a simplified method for Agrobacterium-mediated transformation of *Arabidopsis thaliana*. Plant J 16: 735–743

Conn VM, Hugouvieux V, Nayak A, Conos SA, Capovilla G, Cildir G, Jourdain A, Tergaonkar V, Schmid M, Zubieta C & Conn SJ (2017) A circRNA from SEPALLATA3 regulates splicing of its cognate mRNA through R-loop formation. Nat. plants 3: 17053

Crespi MD, Jurkevitch E, Poiret M, d’Aubenton-Carafa Y, Petrovics G, Kondorosi E & Kondorosi A (1994) enod40, a gene expressed during nodule organogenesis, codes for a non-translatable RNA involved in plant growth. EMBO J. 13: 5099–5112

Ding F, Cui P, Wang Z, Zhang S, Ali S & Xiong L (2014) Genome-wide analysis of alternative splicing of pre-mRNA under salt stress in Arabidopsis. BMC Genomics 15: 431

Djebali S, Davis CA, Merkel A, Dobin A, Lassmann T, Mortazavi A, Tanzer A, Lagarde J, Lin W, Schlesinger F, Xue C, Marinov GK, Khatun J, Williams BA, Zaleski C, Rozowsky J, Roder M, Kokocinski F, Abdelhamid RF, Alioto T, et al (2012) Landscape of transcription in human cells. Nature 489: 101–108

Drechsel G, Kahles A, Kesarwani AK, Stauffer E, Behr J, Drewe P, Ratsch G & Wachter A (2013) Nonsense-mediated decay of alternative precursor mRNA splicing variants is a major determinant of the Arabidopsis steady state transcriptome. Plant Cell 25: 3726–3742

Elvira-Matelot E, Bardou F, Ariel F, Jauvion V, Bouteiller N, Le Masson I, Cao J, Crespi MD & Vaucheret H (2016) The Nuclear Ribonucleoprotein SmD1 Interplays with Splicing, RNA Quality Control, and Posttranscriptional Gene Silencing in Arabidopsis. Plant Cell 28: 426–438

Faial T (2015) RNA splicing in common disease. Nat. Genet. 47: 105

Filichkin SA, Priest HD, Givan SA, Shen R, Bryant DW, Fox SE, Wong W-K & Mockler TC (2010) Genome-wide mapping of alternative splicing in Arabidopsis thaliana. Genome Res. 20: 45–58

Gagnot S, Tamby JP, Martin-Magniette ML, Bitton F, Taconnat L, Balzergue S, Aubourg S, Renou JP, Lecharny A & Brunaud V (2008) CATdb: a public access to Arabidopsis transcriptome data from the URGV-CATMA platform. Nucleic Acids Res. 36: D986–90

Gendrel A-V, Lippman Z, Martienssen R & Colot V (2005) Profiling histone modification patterns in plants using genomic tiling microarrays. Nat. Methods 2: 213–218

Gerstein MB, Rozowsky J, Yan K-K, Wang D, Cheng C, Brown JB, Davis CA, Hillier L, Sisu C, Li JJ, Pei B, Harmanci AO, Duff MO, Djebali S, Alexander RP, Alver BH, Auerbach R, Bell K, Bickel PJ, Boeck ME, et al (2014) Comparative analysis of the transcriptome across distant species. Nature 512: 445–448

Gomez-Gomez L, Felix G & Boller T (1999) A single locus determines sensitivity to bacterial flagellin in Arabidopsis thaliana. Plant J. 18: 277–284

Gonzalez S, Garcia A, Vazquez E, Serrano R, Sanchez M, Quintales L & Antequera F (2016) Nucleosomal signatures impose nucleosome positioning in coding and noncoding sequences in the genome. Genome Res. 26: 1532–1543

Grainger RJ & Beggs JD (2005) Prp8 protein: at the heart of the spliceosome. RNA 11: 533–557

Gultyaev AP & Roussis A (2007) Identification of conserved secondary structures and expansion segments in enod40 RNAs reveals new enod40 homologues in plants. Nucleic Acids Res. 35: 3144–3152

Herold N, Will CL, Wolf E, Kastner B, Urlaub H & Luhrmann R (2009) Conservation of the protein composition and electron microscopy structure of Drosophila melanogaster and human spliceosomal complexes. Mol. Cell. Biol. 29: 281–301

Hutchinson JN, Ensminger AW, Clemson CM, Lynch CR, Lawrence JB & Chess A (2007) A screen for nuclear transcripts identifies two linked noncoding RNAs associated with SC35 splicing domains. BMC Genomics 8: 39

Jabre I, Reddy ASN, Kalyna M, Chaudhary S, Khokhar W, Byrne LJ, Wilson CM & Syed NH (2019) Does co-transcriptional regulation of alternative splicing mediate plant stress responses? Nucleic Acids Res. 47: 2716–2726

Ji Q, Zhang L, Liu X, Zhou L, Wang W, Han Z, Sui H, Tang Y, Wang Y, Liu N, Ren J, Hou F & Li Q (2014) Long non-coding RNA MALAT1 promotes tumour growth and metastasis in colorectal cancer through binding to SFPQ and releasing oncogene PTBP2 from SFPQ/PTBP2 complex. Br. J. Cancer 111: 736–748

Kalyna M, Simpson CG, Syed NH, Lewandowska D, Marquez Y, Kusenda B, Marshall J, Fuller J, Cardle L, McNicol J, Dinh HQ, Barta A & Brown JWS (2012) Alternative splicing and nonsense-mediated decay modulate expression of important regulatory genes in Arabidopsis. Nucleic Acids Res. 40: 2454–2469

Kissen R, Hyldbakk E, Wang C-W V, Sørmo CG, Rossiter JT & Bones AM (2012) Ecotype dependent expression and alternative splicing of epithiospecifier protein (ESP) in Arabidopsis thaliana. Plant Mol. Biol. 78: 361–375

Koncz C, Dejong F, Villacorta N, Szakonyi D & Koncz Z (2012) The spliceosome-activating complex: molecular mechanisms underlying the function of a pleiotropic regulator. Front. Plant Sci. 3: 9

Kong J, Sun W, Li C, Wan L, Wang S, Wu Y, Xu E, Zhang H & Lai M (2016) Long non-coding RNA LINC01133 inhibits epithelial-mesenchymal transition and metastasis in colorectal cancer by interacting with SRSF6. Cancer Lett. 380: 476–484

Kopylova E, Noé L & Touzet H (2012) SortMeRNA: fast and accurate filtering of ribosomal RNAs in metatranscriptomic data. Bioinformatics 28: 3211–3217

Laloum T, Martin G & Duque P (2018) Alternative Splicing Control of Abiotic Stress Responses. Trends Plant Sci. 23: 140–150

Latrasse D, Jégu T, Meng PH, Mazubert C, Hudik E, Delarue M, Charon C, Crespi M, Hirt H, Raynaud C, Bergounioux C & Benhamed M. (2013) Dual function of MIPS1 as a metabolic enzyme and transcriptional regulator. Nucleic Acids Res. 41: 2907–2917

Li Y, Sawada Y, Hirai A, Sato M, Kuwahara A, Yan X & Hirai MY (2013) Novel insights into the function of Arabidopsis R2R3-MYB transcription factors regulating aliphatic glucosinolate biosynthesis. Plant Cell Physiol. 54: 1335–1344

Li Y, Wu Z, Yuan J, Sun L, Lin L, Huang N, Bin J, Liao Y & Liao W (2017) Long non-coding RNA MALAT1 promotes gastric cancer tumorigenicity and metastasis by regulating vasculogenic mimicry and angiogenesis. Cancer Lett. 395: 31–44

Libault M, Wan J, Czechowski T, Udvardi M & Stacey G (2007) Identification of 118 Arabidopsis transcription factor and 30 ubiquitin-ligase genes responding to chitin, a plant-defense elicitor. Mol. Plant. Microbe. Interact. 20: 900–911

Liu S, Kracher B, Ziegler J, Birkenbihl RP & Somssich IE (2015) Negative regulation of ABA signaling by WRKY33 is critical for Arabidopsis immunity towards Botrytis cinerea 2100. Elife 4: e07295

Malakar P, Shilo A, Mogilevsky A, Stein I, Pikarsky E, Nevo Y, Benyamini H, Elgavish S, Zong X, Prasanth K V & Karni R (2017) Long Noncoding RNA MALAT1 Promotes Hepatocellular Carcinoma Development by SRSF1 Upregulation and mTOR Activation. Cancer Res. 77: 1155–1167

Malamy JE & Benfey PN (1997) Organization and cell differentiation in lateral roots of Arabidopsis thaliana. Development 124: 33–44

Marquardt S, Raitskin O, Wu Z, Liu F, Sun Q & Dean C (2014) Functional consequences of splicing of the antisense transcript COOLAIR on FLC transcription. Mol. Cell 54: 156-165

Marquez Y, Brown JWS, Simpson C, Barta A & Kalyna M (2012) Transcriptome survey reveals increased complexity of the alternative splicing landscape in Arabidopsis. Genome Res. 22: 1184–1195

Mercer TR, Dinger ME, Sunkin SM, Mehler MF & Mattick JS (2008) Specific expression of long noncoding RNAs in the mouse brain. Proc Natl Acad Sci USA 105: 716–721

Mercer TR, Qureshi IA, Gokhan S, Dinger ME, Li G, Mattick JS & Mehler MF (2010) Long noncoding RNAs in neuronal-glial fate specification and oligodendrocyte lineage maturation. BMC Neurosci. 11: 14

Moffat CS, Ingle RA, Wathugala DL, Saunders NJ, Knight H & Knight MR (2012) ERF5 and ERF6 play redundant roles as positive regulators of JA/Et-mediated defense against Botrytis cinerea in Arabidopsis. PLoS One 7: e35995

Mohr TJ, Mammarella ND, Hoff T, Woffenden BJ, Jelesko JG & McDowell JM (2010) The Arabidopsis downy mildew resistance gene RPP8 is induced by pathogens and salicylic acid and is regulated by W box cis elements. Mol. Plant. Microbe. Interact. 23: 1303–1315

Munekage YN, Inoue S, Yoneda Y & Yokota A (2015) Distinct palisade tissue development processes promoted by leaf autonomous signalling and long-distance signalling in Arabidopsis thaliana. Plant. Cell Environ. 38: 1116–1126

Musielak TJ, Schenkel L, Kolb M, Henschen A & Bayer M (2015) A simple and versatile cell wall staining protocol to study plant reproduction. Plant Reprod. 28: 161–169

O’Malley RC, Huang S-SC, Song L, Lewsey MG, Bartlett A, Nery JR, Galli M, Gallavotti A & Ecker JR (2016) Cistrome and Epicistrome Features Shape the Regulatory DNA Landscape. Cell 165: 1280–1292

Palusa SG, Ali GS & Reddy ASN (2007) Alternative splicing of pre-mRNAs of Arabidopsis serine/arginine-rich proteins: regulation by hormones and stresses. Plant J. 49: 1091­1107

Pan Q, Shai O, Lee LJ, Frey BJ & Blencowe BJ (2008) Deep surveying of alternative splicing complexity in the human transcriptome by high-throughput sequencing. Nat. Genet. 40: 1413–1415

Pasternak T, Tietz O, Rapp K, Begheldo M, Nitschke R, Ruperti B & Palme K (2015) Protocol: an improved and universal procedure for whole-mount immunolocalization in plants. Plant Methods 11: 50

Pauwels L, Morreel K, De Witte E, Lammertyn F, Van Montagu M, Boerjan W, Inze D & Goossens A (2008) Mapping methyl jasmonate-mediated transcriptional reprogramming of metabolism and cell cycle progression in cultured Arabidopsis cells. Proc. Natl. Acad. Sci. U. S. A. 105: 1380–1385

Pedroza-Garcia J-A, Mazubert C, Del Olmo I, Bourge M, Domenichini S, Bounon R, Tariq Z, Delannoy E, Pineiro M, Jarillo JA, Bergounioux C, Benhamed M & Raynaud C (2017) Function of the Plant DNA Polymerase Epsilon in Replicative Stress Sensing, a Genetic Analysis. Plant Physiol. 173: 1735–1749

Poncini L, Wyrsch I, Denervaud Tendon V, Vorley T, Boller T, Geldner N, Metraux J-P & Lehmann S (2017) In roots of Arabidopsis thaliana, the damage-associated molecular pattern AtPep1 is a stronger elicitor of immune signalling than flg22 or the chitin heptamer. PLoS One 12: e0185808

Pound MP, French AP, Atkinson JA, Wells DM, Bennett MJ & Pridmore T (2013) RootNav: navigating images of complex root architectures. Plant Physiol. 162: 1802–1814

Rapicavoli NA & Blackshaw S (2009) New meaning in the message: noncoding RNAs and their role in retinal development. Dev. Dyn. 238: 2103–2114

Rapicavoli NA, Poth EM & Blackshaw S (2010) The long noncoding RNA RNCR2 directs mouse retinal cell specification. BMC Dev. Biol. 10: 49

Reddy ASN, Marquez Y, Kalyna M & Barta A (2013) Complexity of the alternative splicing landscape in plants. Plant Cell 25: 3657–3683

Rigo R, Bazin J, Crespi M & Charon C (2019) Alternative splicing in the regulation of plant-microbe interactions. Plant Cell Physiol. 60: 1906–1916

Romero-Barrios N, Legascue MF, Benhamed M, Ariel F & Crespi M (2018) Splicing regulation by long noncoding RNAs. Nucleic Acids Res. 46: 2169–2184

Sasaki T, Kanno T, Liang SC, Chen AY, Liao WW, Lin WD, Matzke A & Matzke M (2015) An Rtf2 domain-containing protein influences pre-mRNA splicing and is essential for embryonic development in *Arabidopsis thaliana*. Genetics 200: 523–535

Schenk PM, Kazan K, Rusu AG, Manners JM & Maclean DJ (2005) The *SEN1* gene of Arabidopsis is regulated by signals that link plant defence responses and senescence. Plant Physiol. Biochem. 43: 997–1005

Seo JS, Sun HX, Park BS, Huang CH, Yeh SD, Jung C & Chua NH (2017) ELF18-INDUCED LONG-NONCODING RNA associates with mediator to enhance expression of innate immune response genes in Arabidopsis. Plant Cell 29:1024–1038

Sone M, Hayashi T, Tarui H, Agata K, Takeichi M & Nakagawa S (2007) The mRNA-like noncoding RNA Gomafu constitutes a novel nuclear domain in a subset of neurons. J. Cell Sci. 120: 2498–2506

Song X, Liu G, Huang Z, Duan W, Tan H, Li Y & Hou X (2016) Temperature expression patterns of genes and their coexpression with LncRNAs revealed by RNA-Seq in non-heading Chinese cabbage. BMC Genomics 17: 297

Suresh PS, Tsutsumi R & Venkatesh T (2018) YBX1 at the crossroads of non-coding transcriptome, exosomal, and cytoplasmic granular signaling. Eur. J. Cell Biol. 97: 163–167

Straus MR, Rietz S, Ver Loren van Themaat E, Bartsch M & Parker JE (2010) Salicylic acid antagonism of EDS1-driven cell death is important for immune and oxidative stress responses in Arabidopsis. Plant J. 62: 628–640

Syed NH, Kalyna M, Marquez Y, Barta A & Brown JWS (2012) Alternative splicing in plants--coming of age. Trends Plant Sci. 17: 616–623

Tanabe N, Yoshimura K, Kimura A, Yabuta Y & Shigeoka S (2007) Differential expression of alternatively spliced mRNAs of Arabidopsis SR protein homologs, atSR30 and atSR45a, in response to environmental stress. Plant Cell Physiol. 48: 1036–1049

Tanackovic G, Ransijn A, Ayuso C, Harper S, Berson EL & Rivolta C (2011) A missense mutation in PRPF6 causes impairment of pre-mRNA splicing and autosomal-dominant retinitis pigmentosa. Am. J. Hum. Genet. 88: 643–649

Tran VDT, Souiai O, Romero-Barrios N, Crespi M & Gautheret D (2016) Detection of generic differential RNA processing events from RNA-seq data. RNA Biol. 13: 59–67

Trincado JL, Entizne JC, Hysenaj G, Singh B, Skalic M, Elliott DJ & Eyras E (2018) SUPPA2: fast, accurate, and uncertainty-aware differential splicing analysis across multiple conditions. Genome Biol. 19: 40

Tsuiji H, Yoshimoto R, Hasegawa Y, Furuno M, Yoshida M & Nakagawa S (2011) Competition between a noncoding exon and introns: Gomafu contains tandem UACUAAC repeats and associates with splicing factor-1. Genes Cells 16: 479–490

van Dijk M, Visser A, Buabeng KML, Poutsma A, van der Schors RC & Oudejans CBM (2015) Mutations within the LINC-HELLP non-coding RNA differentially bind ribosomal and RNA splicing complexes and negatively affect trophoblast differentiation. Hum. Mol. Genet. 24: 5475–5485

Villamizar O, Chambers CB, Riberdy JM, Persons DA & Wilber A (2016) Long noncoding RNA Saf and splicing factor 45 increase soluble Fas and resistance to apoptosis. Oncotarget 7: 13810–13826

Vitting-Seerup K & Sandelin A (2017) The landscape of isoform switches in human cancers. Mol. Cancer Res. 15: 1206–1220

Vitting-Seerup K & Sandelin A (2019) IsoformSwitchAnalyzeR: analysis of changes in genome-wide patterns of alternative splicing and its functional consequences. Bioinformatics 35: 4469–4471

Wang D, Ding L, Wang L, Zhao Y, Sun Z, Karnes RJ, Zhang J & Huang H (2015) LncRNA MALAT1 enhances oncogenic activities of EZH2 in castration-resistant prostate cancer. Oncotarget 6: 41045–41055

Wang ET, Sandberg R, Luo S, Khrebtukova I, Zhang L, Mayr C, Kingsmore SF, Schroth GP & Burge CB (2008) Alternative isoform regulation in human tissue transcriptomes. Nature 456: 470–476

Wang H, Wang J, Jiang J, Chen S, Guan Z, Liao Y & Chen F (2014) Reference genes for normalizing transcription in diploid and tetraploid Arabidopsis. Sci. Rep. 4: 6781

West JA, Davis CP, Sunwoo H, Simon MD, Sadreyev RI, Wang PI, Tolstorukov MY & Kingston RE (2014) The long noncoding RNAs NEAT1 and MALAT1 bind active chromatin sites. Mol. Cell 55: 791–802

Whittaker C & Dean C (2017) The FLC Locus: A Platform for Discoveries in Epigenetics and Adaptation. Annu. Rev. Cell Dev. Biol. 33: 555–575

Xu S, Zhang Z, Jing B, Gannon P, Ding J, Xu F, Li X & Zhang Y (2011) Transportin-SR is required for proper splicing of resistance genes and plant immunity. PLoS Genet. 7: e1002159

Yan K, Liu P, Wu C-A, Yang G-D, Xu R, Guo Q-H, Huang J-G & Zheng C-C (2012) Stress-induced alternative splicing provides a mechanism for the regulation of microRNA processing in Arabidopsis thaliana. Mol. Cell 48: 521–531

Yin Q-F, Yang L, Zhang Y, Xiang J-F, Wu Y-W, Carmichael GG & Chen L-L (2012) Long noncoding RNAs with snoRNA ends. Mol. Cell 48: 219–230

Yoshida K, Sanada M, Shiraishi Y, Nowak D, Nagata Y, Yamamoto R, Sato Y, Sato-Otsubo A, Kon A, Nagasaki M, Chalkidis G, Suzuki Y, Shiosaka M, Kawahata R, Yamaguchi T, Otsu M, Obara N, Sakata-Yanagimoto M, Ishiyama K, Mori H, et al (2011) Frequent pathway mutations of splicing machinery in myelodysplasia. Nature 478: 64–69

Zhan X, Qian B, Cao F, Wu W, Yang L, Guan Q, Gu X, Wang P, Okusolubo TA, Dunn SL, Zhu J-K & Zhu J (2015) An Arabidopsis PWI and RRM motif-containing protein is critical for pre-mRNA splicing and ABA responses. Nat. Commun. 6: 8139

Zhang R, Calixto CPG, Marquez Y, Venhuizen P, Tzioutziou NA, Guo W, Spensley M, Entizne JC, Lewandowska D, Ten Have S, Frei Dit Frey N, Hirt H, James AB, Nimmo HG, Barta A, Kalyna M & Brown JWS (2017) A high quality Arabidopsis transcriptome for accurate transcript-level analysis of alternative splicing. Nucleic Acids Res. 45: 5061–5073

Zhang X-N, Mo C, Garrett WM & Cooper B (2014) Phosphothreonine 218 is required for the function of SR45.1 in regulating flower petal development in Arabidopsis. Plant Signal. Behav. 9: e29134

Zhang Y, Zhang X-O, Chen T, Xiang J-F, Yin Q-F, Xing Y-H, Zhu S, Yang L & Chen L-L (2013) Circular intronic long noncoding RNAs. Mol. Cell 51: 792–806

Zhang Z, Liu Y, Ding P, Li Y, Kong Q & Zhang Y (2014) Splicing of receptor-like kinase-encoding SNC4 and CERK1 is regulated by two conserved splicing factors that are required for plant immunity. Mol. Plant 7: 1766–1775

Zipfel C, Robatzek S, Navarro L, Oakeley EJ, Jones JDG, Felix G & Boller T (2004) Bacterial disease resistance in Arabidopsis through flagellin perception. Nature 428: 764–767

