## Supplementary figures and images for "An *Arabidopsis* long noncoding RNA modulates the transcriptome through interactions with a network of splicing factors"

### EV1

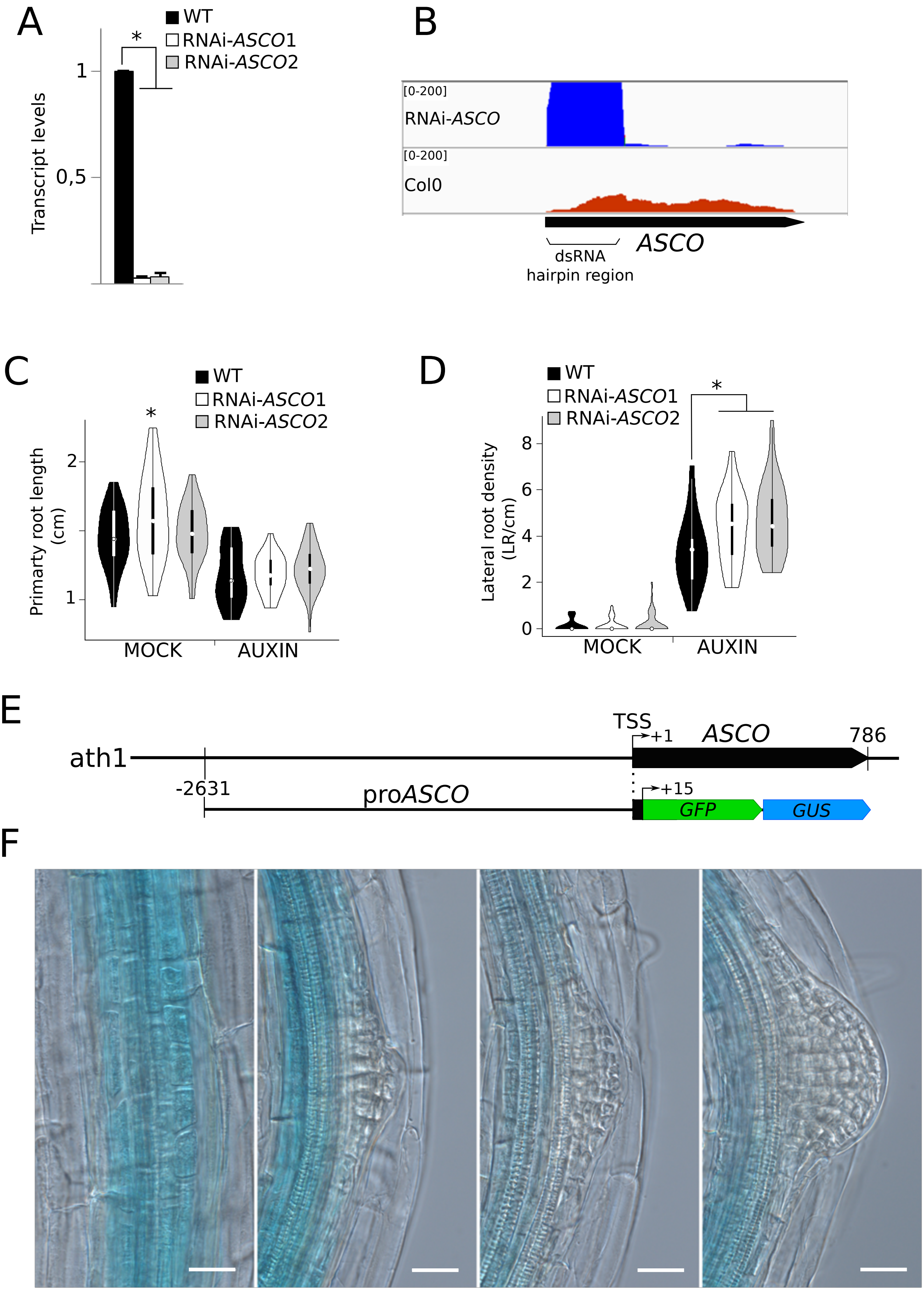

### EV2

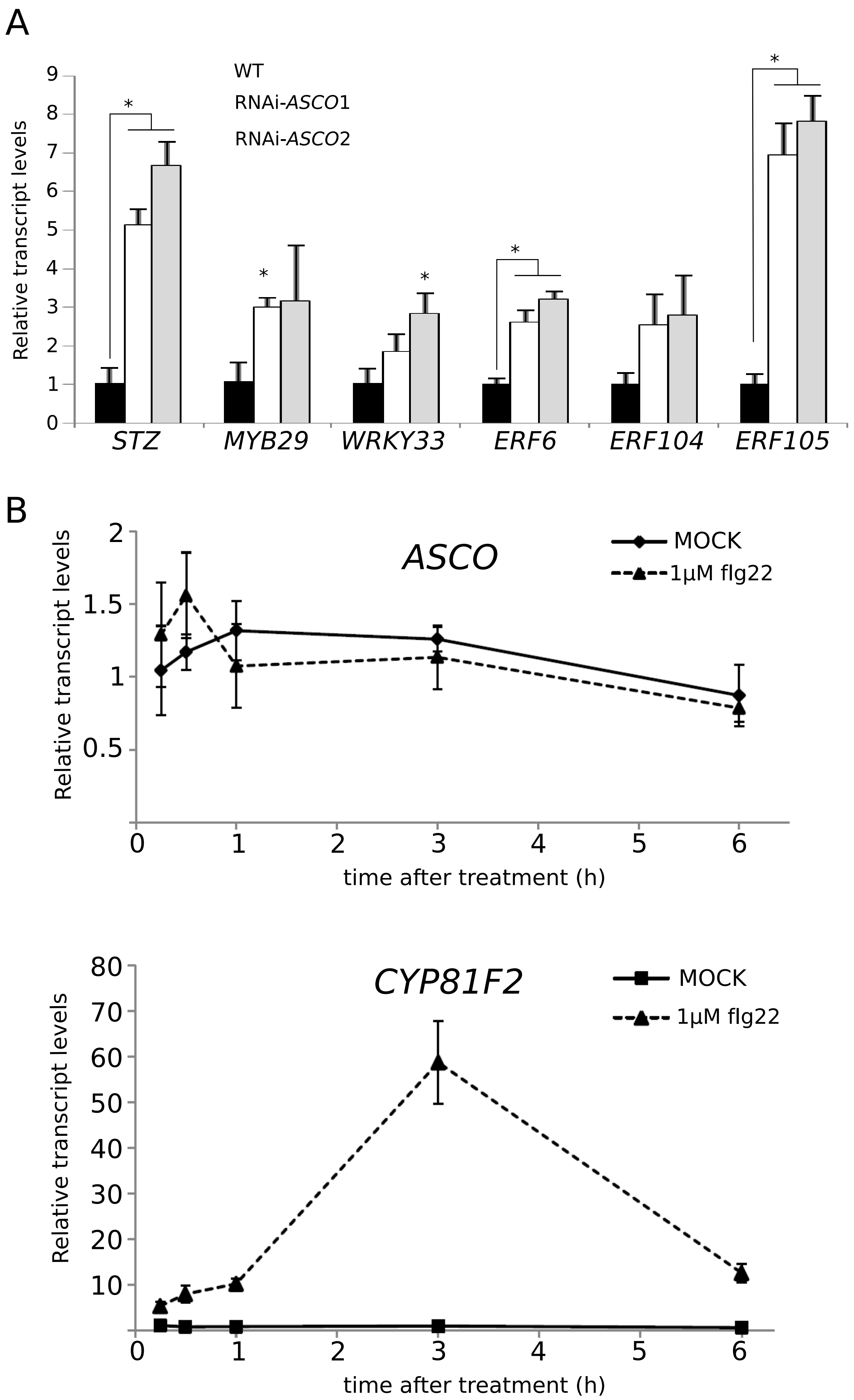

### EV3

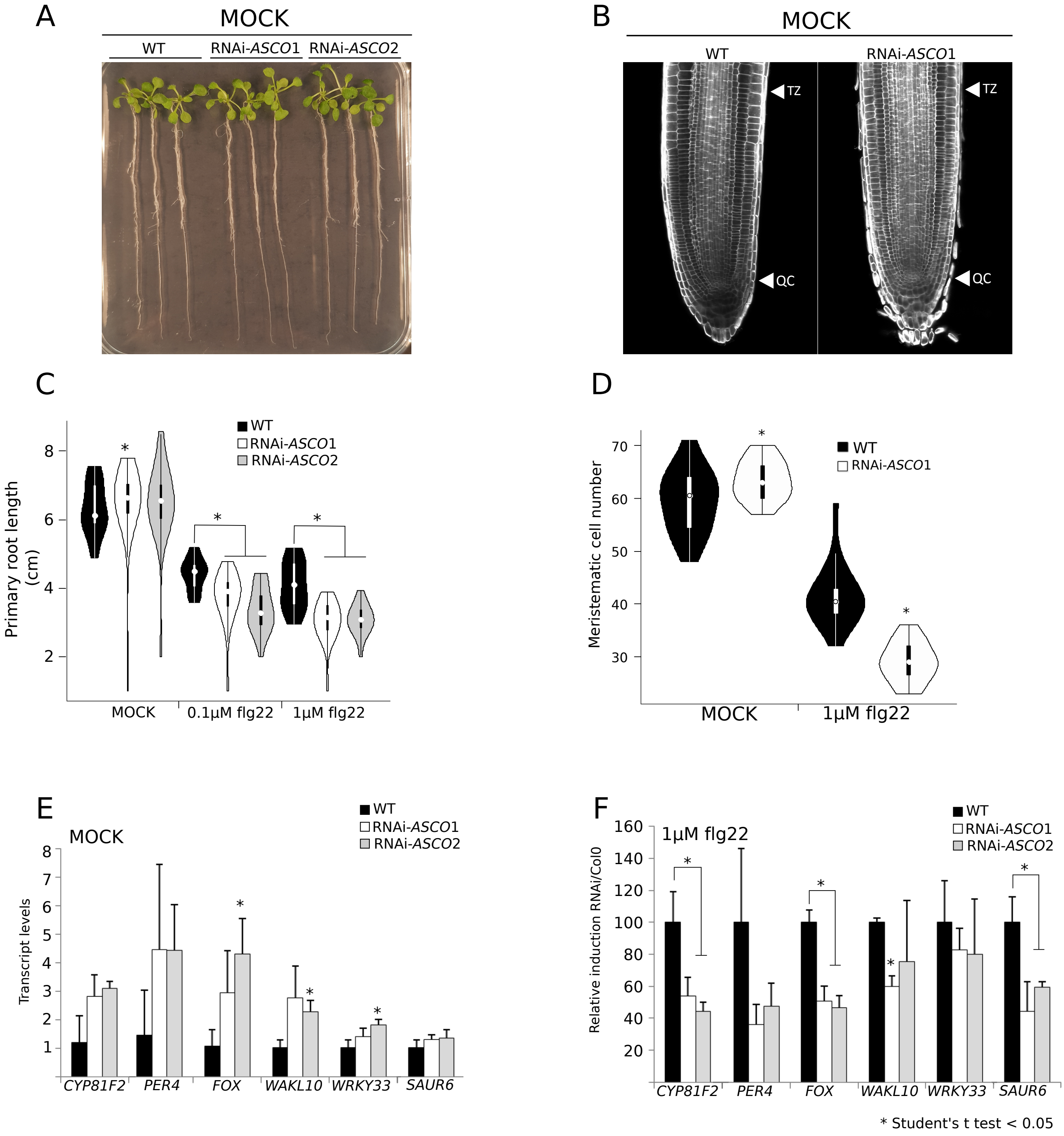

### EV4

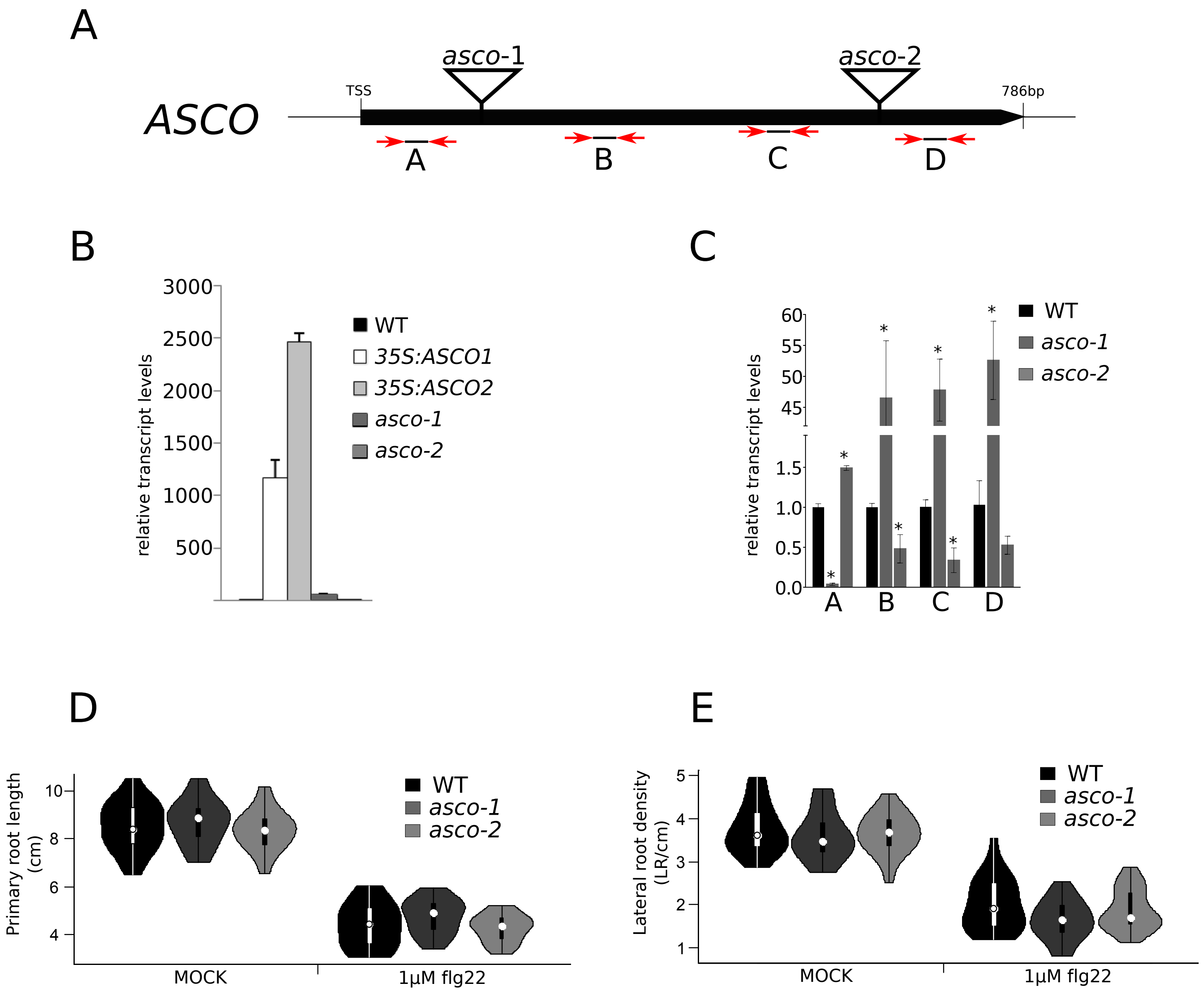

### EV5

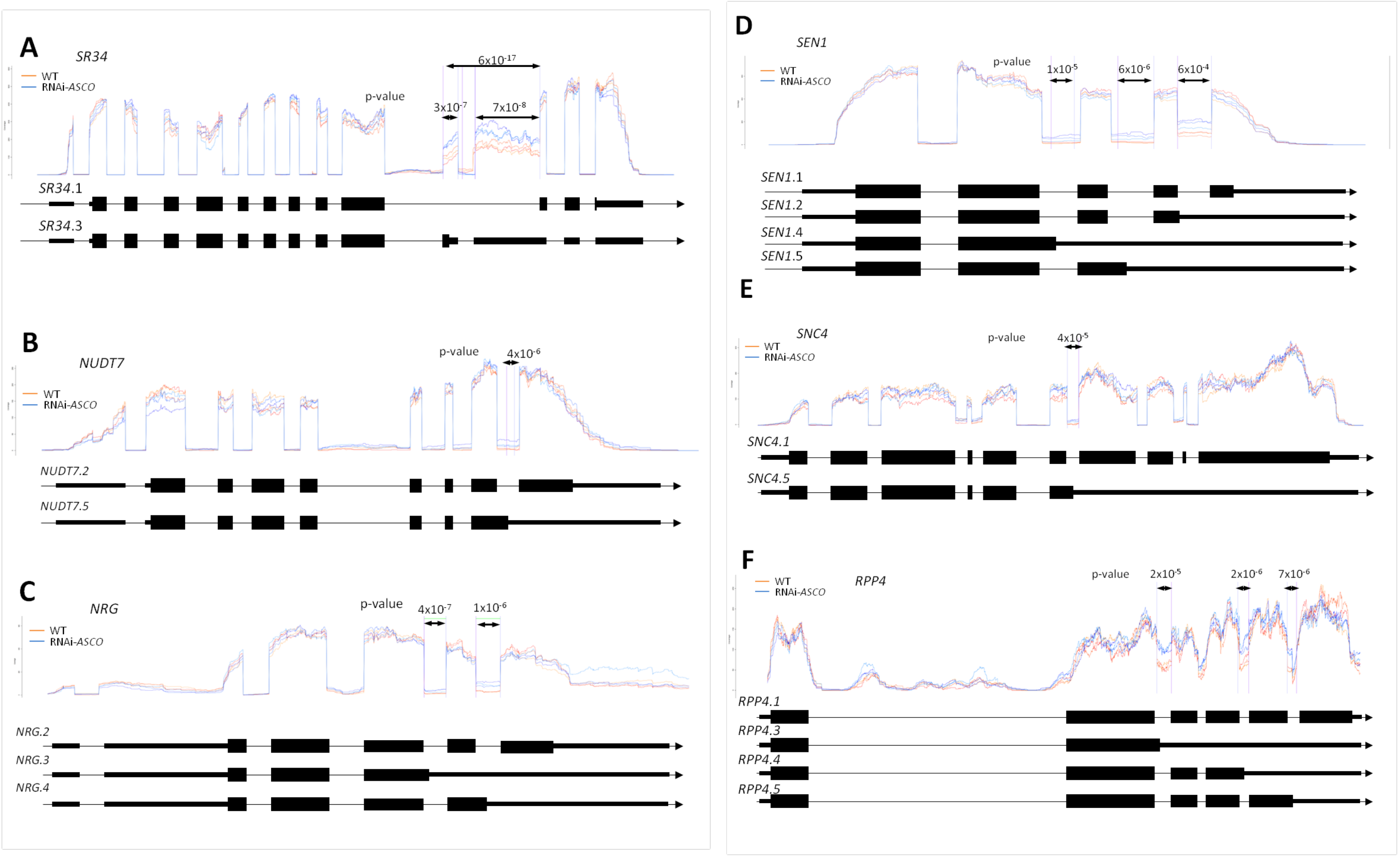

### EV6

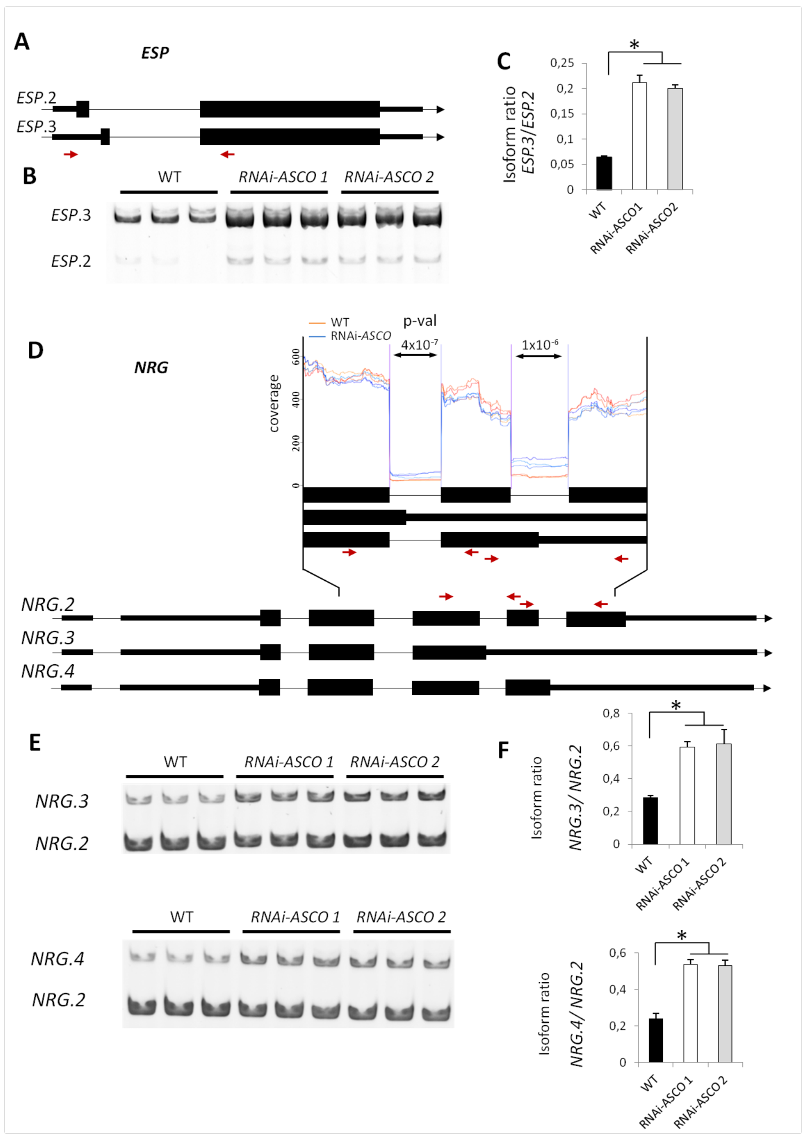

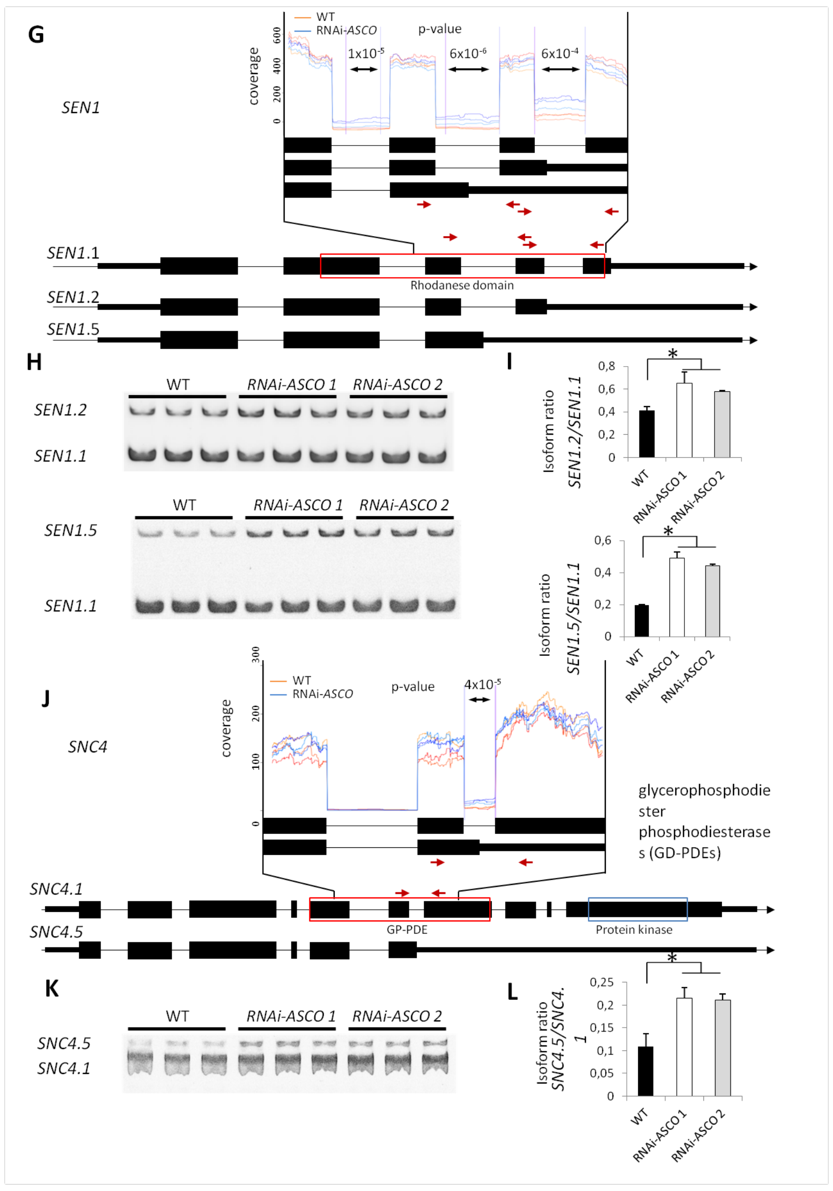

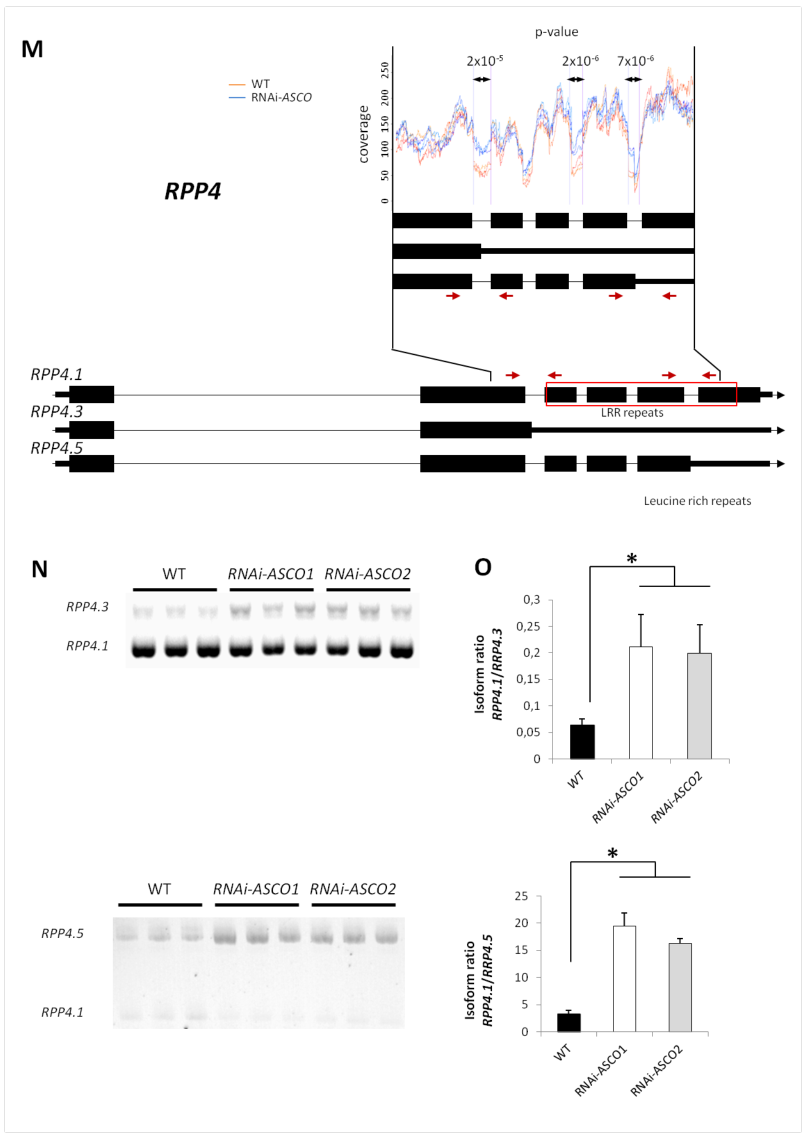

### EV7

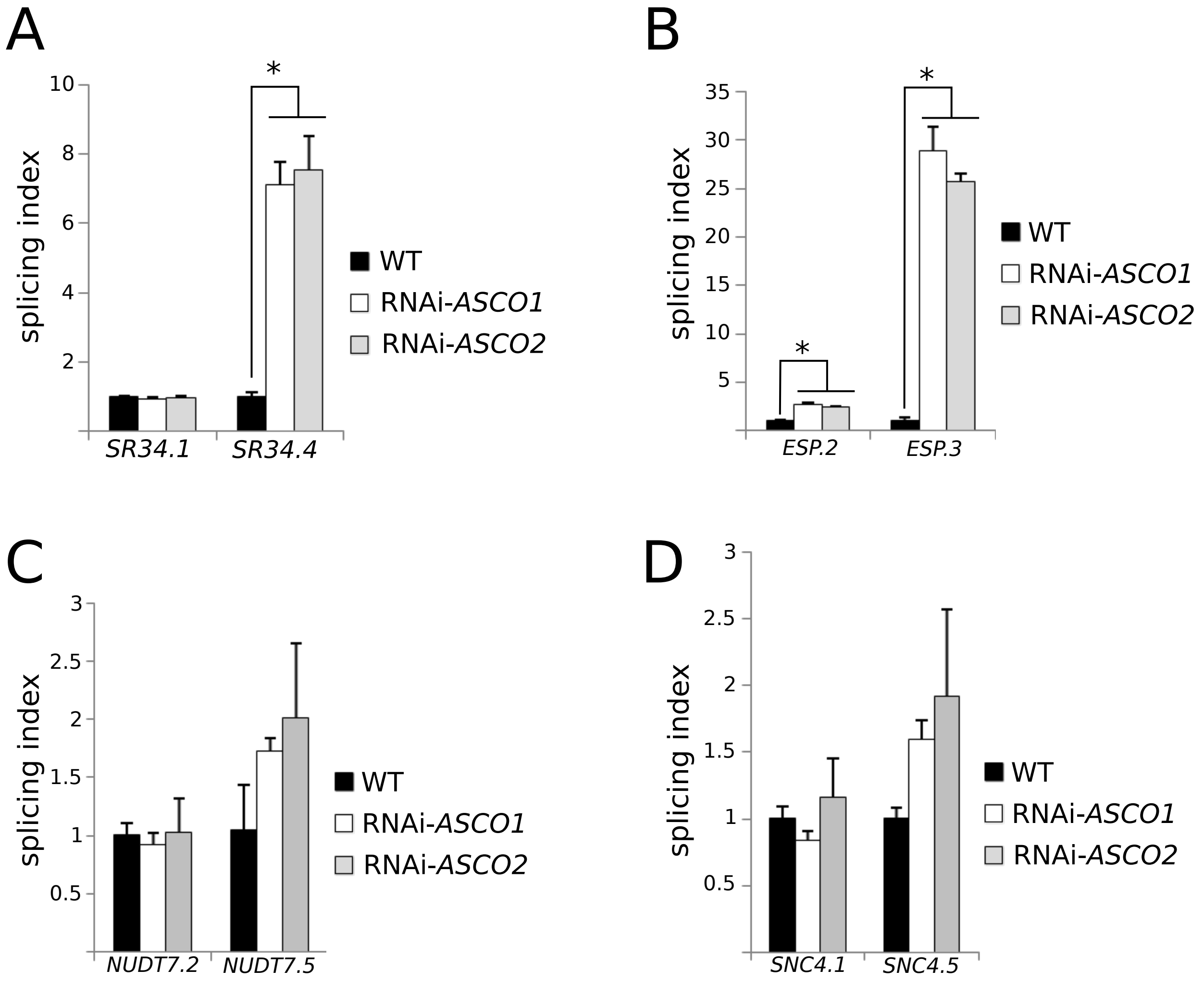

### EV8

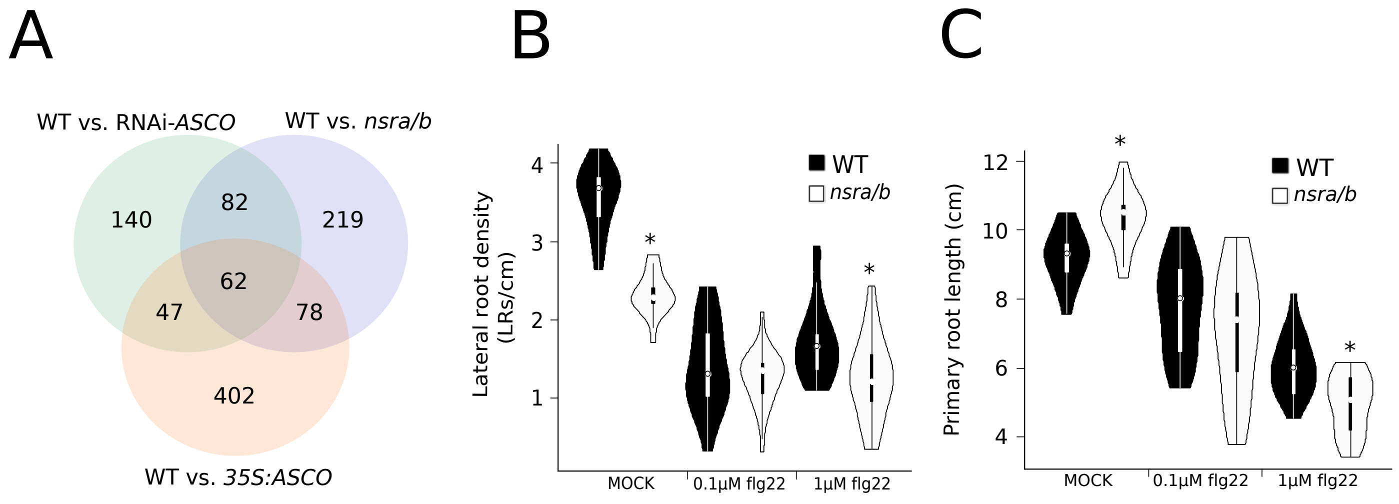

### EV9

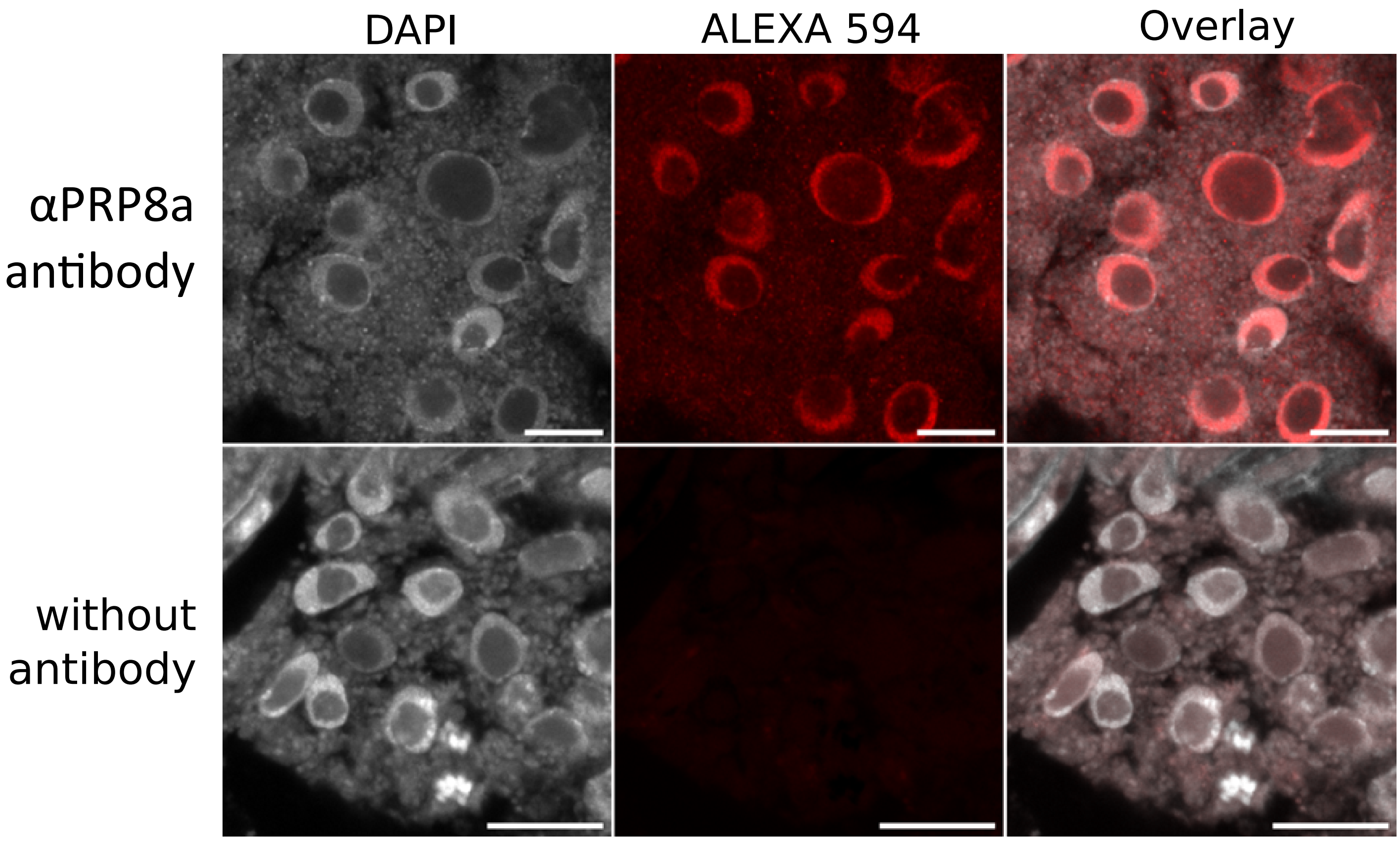

### EV10

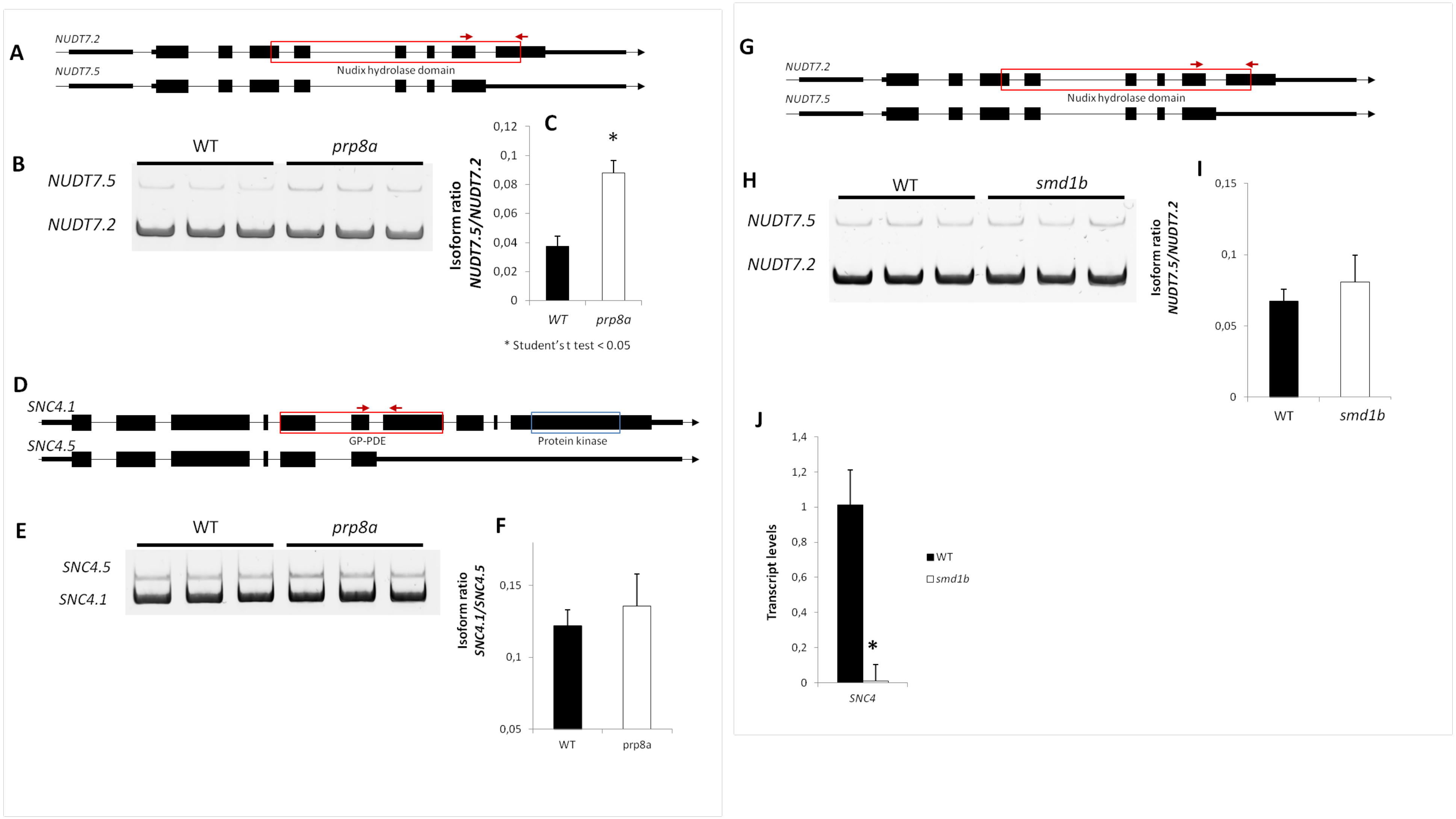

### EV11

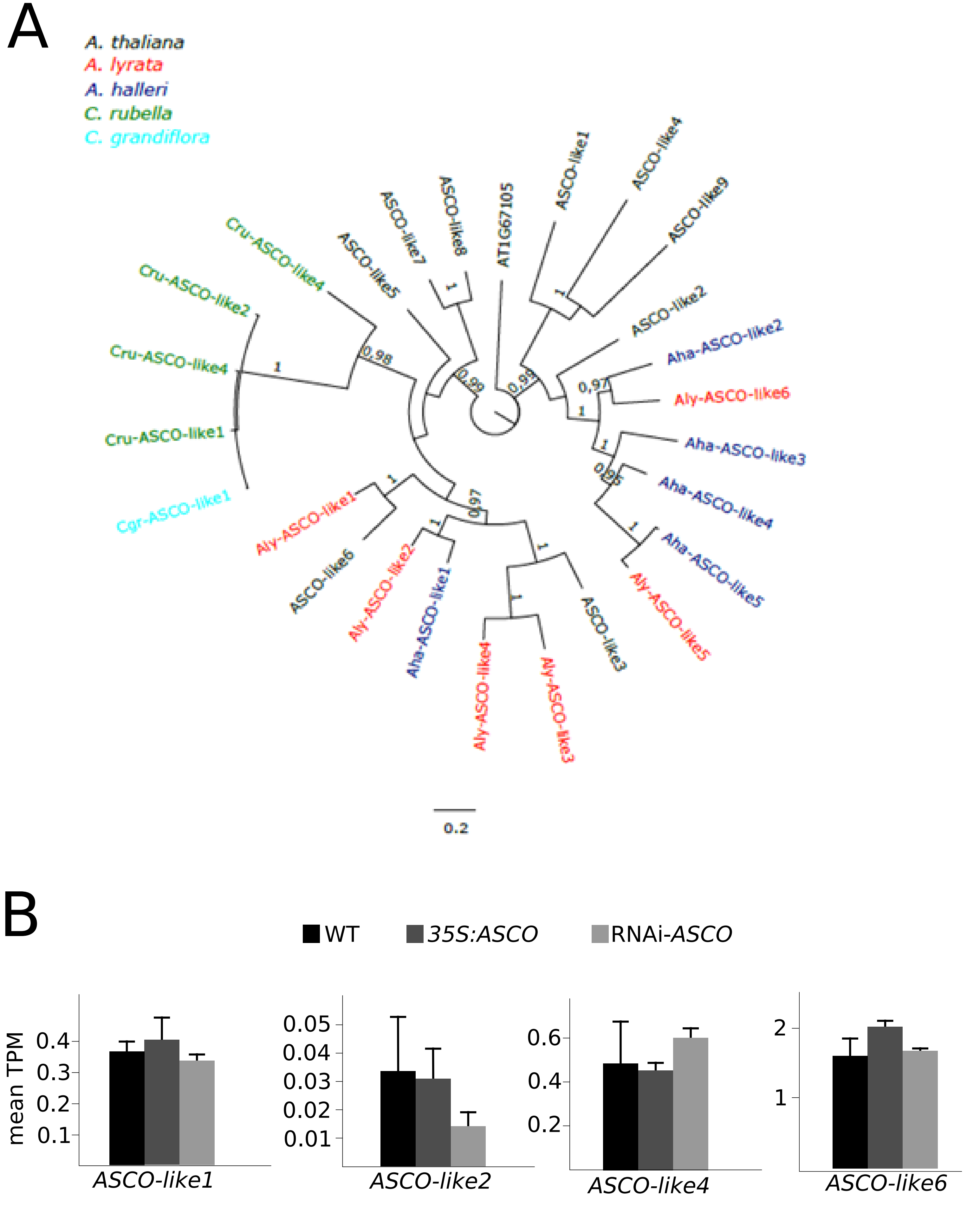
